# Harnessing machine learning models for epigenome to transcriptome association studies

**DOI:** 10.1101/2025.05.09.653095

**Authors:** Fatemeh Behjati Ardakani, Shamim Ashrafiyan, Laura Rumpf, Dennis Hecker, Marcel H Schulz

## Abstract

Understanding how epigenome variation contributes to gene expression in disease and development is a fundamental challenge. Regulatory regions show cell type-specific epigenome activity and differ in their location, size, and distance to their target genes, complicating discovery and analysis. Recent machine learning models have been proposed to address these problems by learning functions for the prediction of gene expression from epigenomic data. Here, we use the large IHEC EpiATLAS dataset to benchmark state-of-the-art linear and non-linear approaches. Each approach is optimized for over 28,000 human genes, providing a comprehensive regulatory catalog of gene models. In-depth comparison reveals that gene characteristics and the epigenomic complexity of the locus influence the difficulty of predicting the epigenome-to-transcriptome association. The model performance is further evaluated using CRISPRi and eQTL validation data. Based on these models, we conduct histone-acetylation association studies in a systematic way to investigate how epigenomic variation impacts gene expression. The model-based analysis revealed genes and regulatory regions linked to B-cell leukemia in patient data with known disease-related functions. Our work provides a foundation for applications that link epigenome variation to gene expression in human cells, by benchmarking methods on a per-gene basis, illustrating their use in a disease context and making trained models available to the community.

## Introduction

Epigenetic regulation is central to basic cellular processes, such as transcription and post-transcriptional regulation, and thus important in human development and disease. Genome-wide measurements of histone marks via ChIP-seq revealed associations with different elements, including gene bodies and cis-regulatory elements (CREs) [1]. Several histone marks are known to associate with active regulatory regions, e.g., the modifications H3K4me3 and H3K27ac, and are therefore used to locate cell type-specific CREs [2, 3]. It has become routine to generate H3K27ac ChIP-seq datasets for case-control medical cohorts to study epigenome variation in disease and identify CREs and genes that are connected to disease in a cell- or tissue-specific way using histone-acetylome-wide association studies (HAWAS) [4–10]. Thus far, a HAWAS is done by first finding H3K27ac peaks that are associated with the disease in question, then other integrative analyses are performed to link peaks to genes, which are then considered disease-related genes. Despite the availability of large datasets of histone marks from disease cohorts, there are few specific methods for such association studies, although progress has been made for methods that allow to associate epigenome variation with changes in gene expression.

In general, there are two different categories of approaches for learning epigenome-expression associations [11]. The first category is to build a gene-agnostic model, which treats all genes as equivalent training instances for associating epigenome and expression data from a specific biological sample for model training [12–19]. The second is to build one model per gene (gene-specific) using many paired samples of epigenome and expression data [20–22]. The former does not depend on the availability of numerous biological samples, as it leverages the multitude of genes within a single genome. However, it has the limitation of being unable to accurately capture gene-specific locations of CREs and their respective strengths of association with gene expression. The latter addresses this caveat, but requires many datasets and is computationally intensive as tens of thousands of models need to be trained. Thus, previous studies have used simple linear models for gene-specific approaches, although there is evidence that modeling nonlinear associations between histone marks and gene expression is useful [18, 19]. This work contributes to the field in the following ways: (i) we compare state-of-the-art machine learning approaches for predicting gene expression from H3K27ac data in a gene-specific manner, (ii) we devise a comprehensive benchmark on the large EpiATLAS dataset from IHEC [23], and (iii) inspired by model-based transcriptome-wide association studies [24], we propose two new principled approaches to conduct a HAWAS using prediction models (Fig. 1). The first approach reveals genes associated with epigenetic variation in the disease. The second approach identifies the regulatory regions linked to disease-relevant genes. We illustrate both types of HAWAS on a dataset from patients with chronic lymphocytic leukemia and suggest novel genes and regions associated with the disease. The best models are made available to the community for further use on other datasets via the EpiExpress repository.

**Fig. 1.**
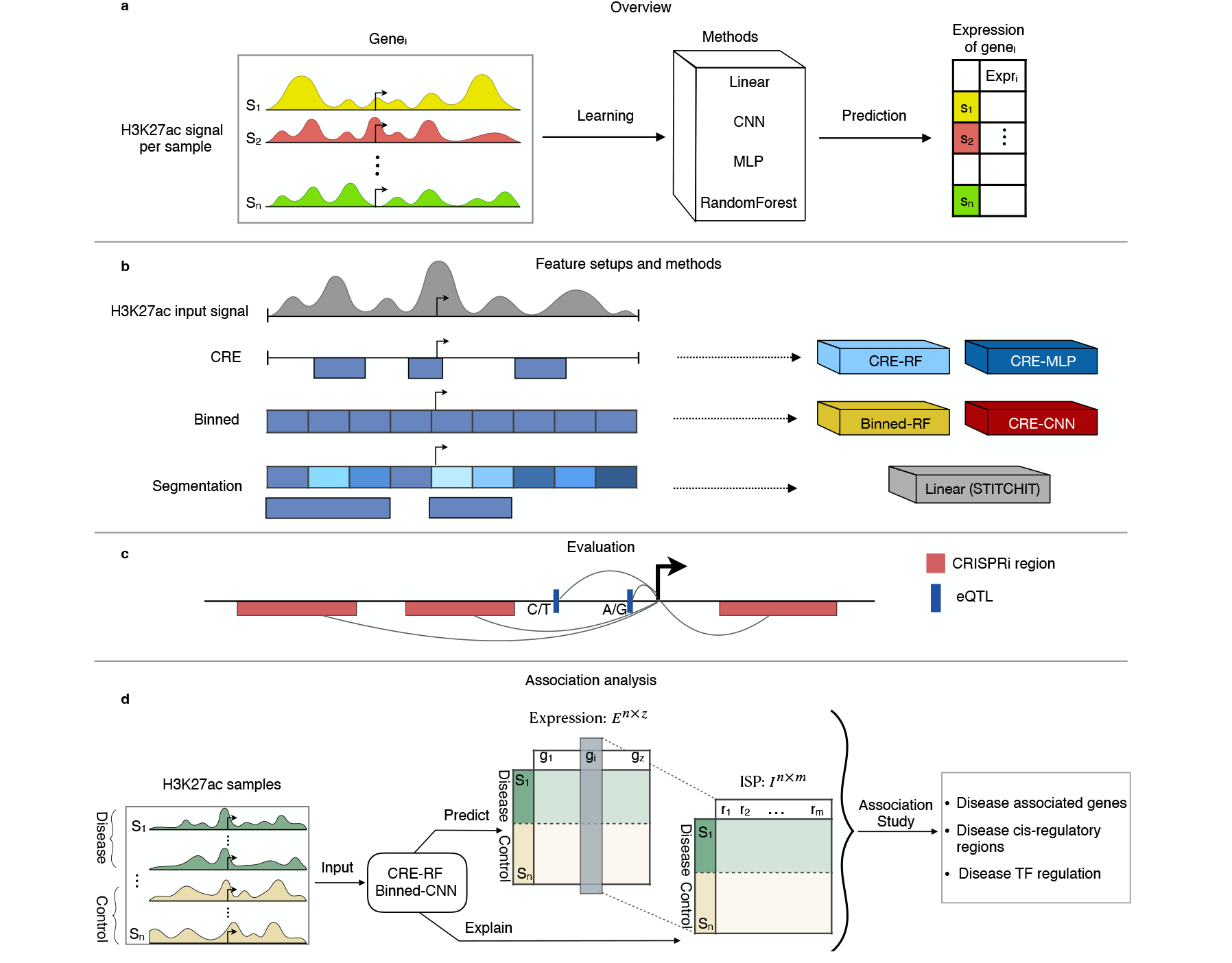
Overview of the model training, feature setups, evaluation of the predicted enhancer-gene interactions and disease-association analysis. **(a)** Each model is learned per gene to predict gene expression using H3K27ac signal for *n* samples around a gene as input. **(b)** H3K27ac signal is quantified for different types of feature setups that are combined with machine learning approaches to form five methods for learning. **(c)** Validation is done using external eQTL and CRISPRi data. **(d)** Overview of the analysis workflow for a histone-acetylome-wide association study using the CRE-RF and Binned-CNN models. Gene expression matrix (*E* ∈ ℝ^*n×z*^), where *n* is the number of samples and *z* is the number of genes, is predicted using either method from H3K27ac signal of control and disease samples. Statistical analysis of the predicted gene expression values is used to identify disease-associated genes. Analysis of *in silico* perturbation values (ISPs, Fig. 4) reveals disease-associated regulatory regions, represented by the matrix *I* ∈ ℝ^*n×m*^, where *m* is the number of regions, as well as the involved transcription factors (TFs).

## Results

### Benchmarking of state-of-the-art machine learning methods for expression prediction

The application of epigenome-to-expression prediction models relies on having accurate models. To evaluate whether current methodology yields such accurate models, we conducted a comprehensive benchmark study for gene-specific approaches (Fig. 1a). The benchmark dataset comprised 965 samples originating from 51 cell types/tissues for which both RNA-seq and H3K27ac ChIP-seq data were available in the IHEC EpiATLAS (https://ihec-epigenomes.org/epiatlas/data/)(Supp. Fig. 1a). We applied state-of-the-art machine learning approaches to the task of gene-specific learning. Non-linear associations were accommodated using convolutional neural network (CNN) and multilayer perceptron (MLP) architectures (Supp. Fig. 2), and random forests (RFs). In addition, we used **STITCHIT**, a method that combines signal segmentation and sparse linear regression [22].

We applied various strategies to generate a collection of putative regulatory elements as features for the subsequent model learning step (Fig. 1b). The different feature setups result in varying numbers of input features (Supp. Fig. 1b). We quantified the epigenetic signal in predefined consensus cis-regulatory elements (CREs) identified by the ENCODE project [25] in a 1 MB genomic window, surrounding the most 5’ GENCODE TSS of a gene (**CRE** strategy). The corresponding models are denoted as *CRE-MLP* and *CRE-RF*. Additionally, we introduced a **binned** approach, where the features were defined as 10,000 consecutive, non-overlapping bins of 100 base pairs (bp) [13], which gave the models full flexibility to weigh regions of interest. This specific feature configuration was chosen for an RF-based approach (*Binned-RF*) and for a CNN-based approach referred to as *Binned-CNN*. The linear **STITCHIT** algorithm incorporates an integrated feature selection approach that starts by dividing the genome into small equal-sized bins, similar to the binned strategy, but successively merges adjacent bins into larger segments. Applying CNNs to the CRE-based or STITCHIT setup is not appropriate, as the CRE regions are not adjacent and are unsuitable for convolutional operations. Thus, in total five different gene-specific machine learning (ML) approaches were compared that differed in their architecture and ability to handle nonlinear associations.

### Comparison reveals gene-specific challenges for accurate prediction

All subsequent analyses were performed on 28, 180 genes that passed our quality filtering criteria on the EpiATLAS. Parameter optimization was done for each gene and each ML approach separately. For evaluation, mean squared error (MSE) and the Pearson correlation coefficient between predicted and actual gene expression were calculated on unseen test samples.

In order to better characterize the resulting gene models, a partitioning into three quality sets (*high* ≥ 0.7, *intermediate* ≥ 0.3 and *fail <* 0.3) based on test correlation was obtained (Fig. 2a). The *fail* class also included cases for which a method did not produce any model. The methods with the largest fractions of models achieving high correlation values were CRE-RF (41%) and Binned-CNN (40%). The CRE-RF was also the method with the lowest percentage of failed models (1%). Across genes, all models showed large variance in their accuracy to predict unseen samples (Supp. Fig. 3a), highlighting the challenge of learning models per gene. The lowest median error was obtained for CRE-RF (MSE 3.8), closely followed by the Binned-CNN (MSE 3.85). Of all the methods, CRE-MLP had the highest error (median MSE of 4.19). For some genes, some methods failed to obtain a reliable model, which was particularly often the case for STITCHIT due to internal filtering on discretized gene expression (failed for 11,547 genes) (Supp. Fig. 3b).

**Fig. 2.**
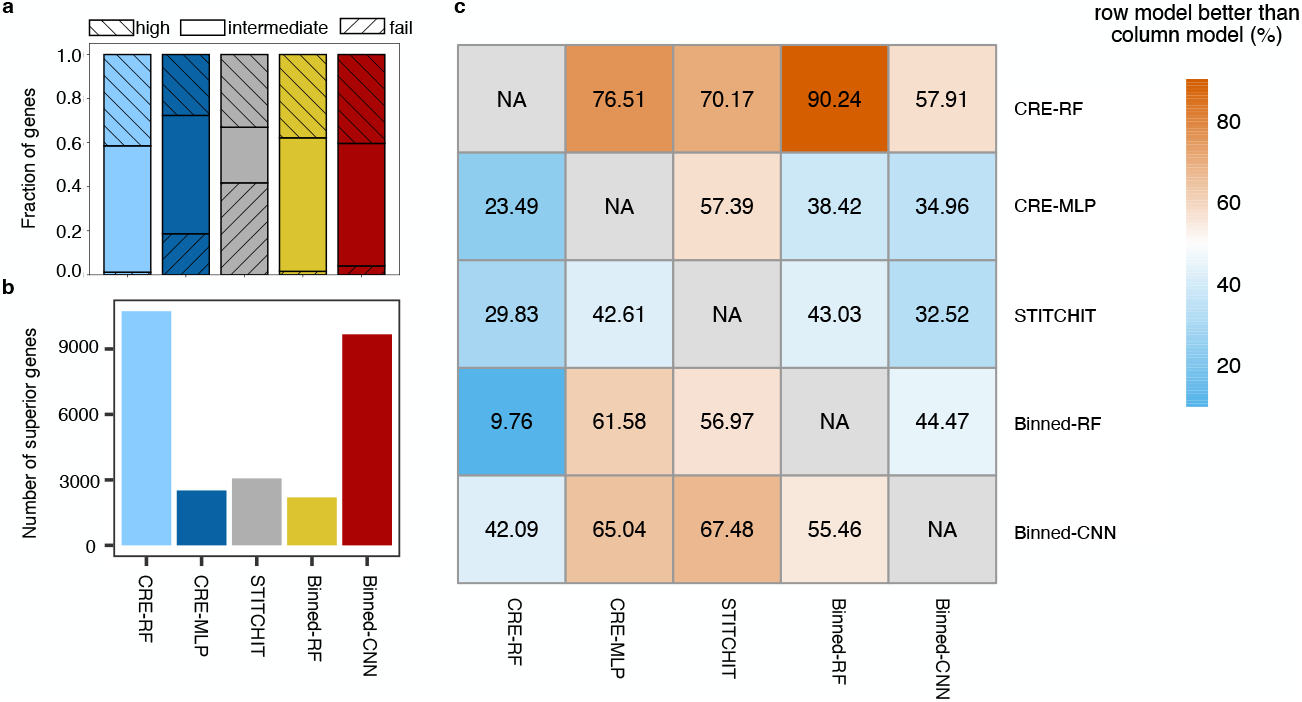
Performance assessment of the five machine learning approaches. **(a)** Genes are partitioned according to performance (Pearson correlation) into the classes *high* (≥ 0.7), *intermediate* (≥ 0.3) and *fail* (< 0.3) for each method. **(b)** Barplot of the number of genes where a method out-performs all other methods (best Pearson correlation). **(c)** Heatmap displaying pairwise comparisons between methods, illustrating the percentage of genes for which the method in each row outperforms the method in each column, based on MSE. Correlation and MSE are always estimated on the test set.

To determine whether any single approach consistently outperformed all other methods, the number of gene models where a given method achieved the highest Pearson correlation was obtained (Fig. 2b). CRE-RF ranked first (11, 404 genes), followed by Binned-CNN (10, 004 genes). However, each method had a subset of genes for which its performance was uniquely superior. For a more detailed analysis, the ML approaches were compared pairwise. Figure 2c shows the percentage of a method (row) outperforming another method (column) based on MSE (other metrics in Supp. Fig. 3c,d). The CRE-based feature strategy outperformed the binned strategy for 90.23% of the genes when comparing the CRE-RF with the Binned-RF, underlining the importance of feature selection in such challenging learning problems. When comparing the two best-performing methods, CRE-RF outperformed Binned-CNN in 57.90% of cases.

To gain insight into what could explain the variable performance of ML approaches in gene subsets, a number of gene-related descriptors were investigated that may affect training or generalization performance. The descriptors were grouped into three categories related to *structure*, e.g., gene length, gene *expression* variability or *genome context*, e.g., number of CREs in the surrounding area (Supp. Fig. 4a) which showed a large variability between genes (Fig. 3a).

**Fig. 3.**
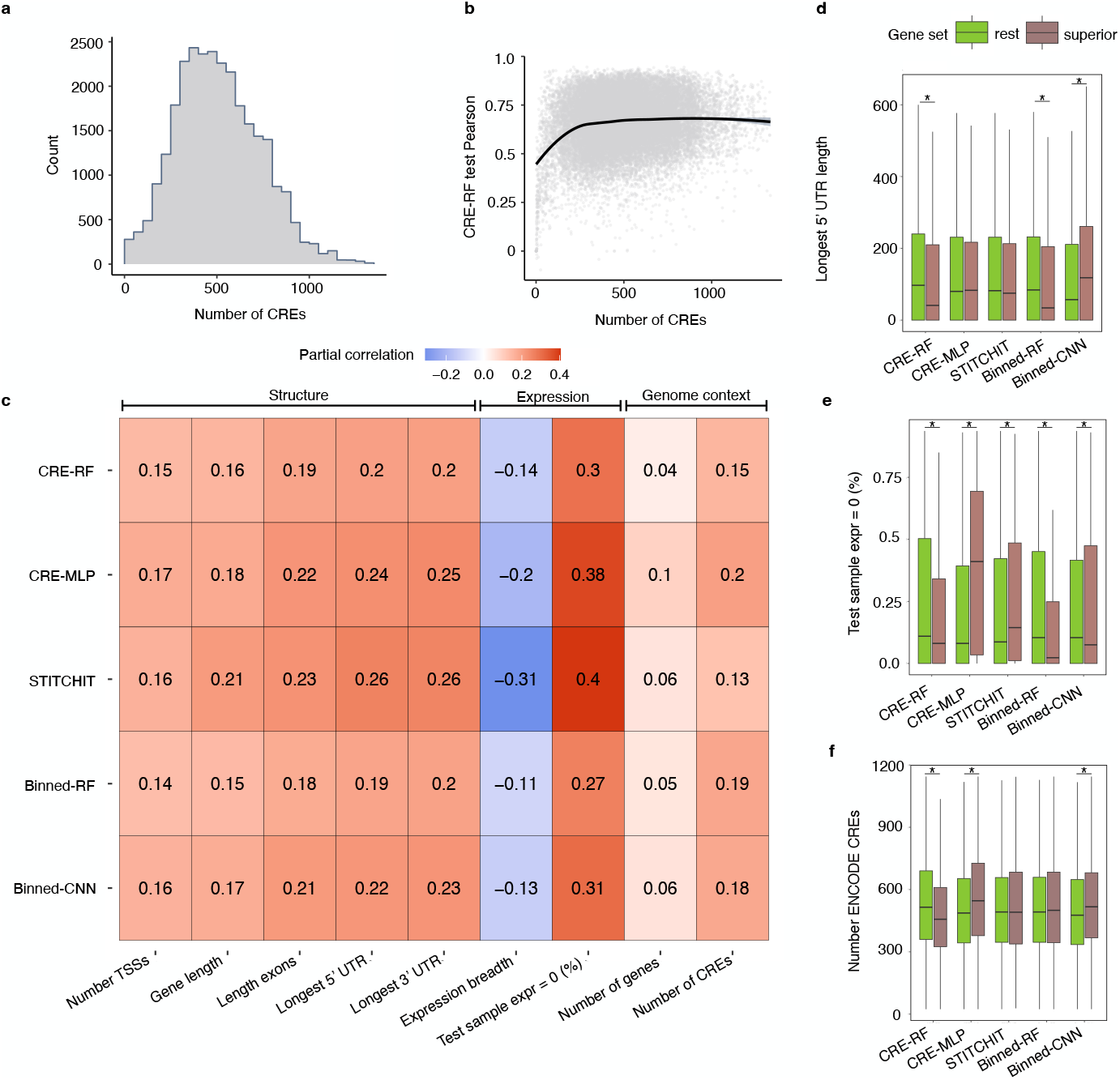
Investigation of gene characteristics that affect model performance. **(a)** Histogram for the number of gene regulatory regions (CREs) located within a 1 MB gene window. **(b)** Scatter-plot comparing the number of CREs and model performance of the CRE-RF (Pearson correlation). **(c)** Different gene descriptors were organized into three categories that characterize a gene’s *structure, expression* and *genome context* (Supp. Fig. 4a). The impact of one descriptor with the model performance (Pearson correlation) was assessed using partial correlation analyses to correct for confounding correlated descriptors from other categories. The Heatmap shows partial correlation values between each model (row) and descriptors grouped by category (columns). Structure: number of TSSs, gene length, total exon length, lengths of the longest 5’ and 3’ UTR; Expression: fraction of cell types/tissues (n = 58) in the entire dataset where the target gene is expressed (TPM ≥ 0.5), fraction of test samples with expression = 0; Genome context: number of other genes or number of CREs in the 1 MB gene window. **(d**,**e**,**f)** Investigation of genes for which a method has the best MSE value (superior) and all remaining genes (rest) for three descriptors. Boxplot for the length of the longest 5’ UTR **(d)**, the fraction of test samples with expression = 0 and **(e)** the number of CREs in the 1 MB window **(f).** Center line is median, boxlimits correspond to IQR, whiskers to 1.5x IQR. Outliers are not shown. An unpaired Mann-Whitney U Test was performed between the superior and rest gene sets for each gene descriptor and each method separately (*p*-value ≤ 0.05 indicated with an asterisk). Supp. Fig. 4d-i show the boxplots for the remaining descriptors. Pearson and partial correlation values were estimated on the test set.

A systematic assessment of all gene descriptors and model accuracy was performed using a partial correlation approach that corrects for performance loss caused by descriptors from other categories (Fig. 3b,c). All structural descriptors had a positive association with model performance for all approaches. For all methods, the strongest positive influence on model performance was observed for the number of test samples with zero expression. This indicates that all methods could predict tissue-specific genes more easily. Accordingly, the negative impact of the expression breadth shows that it was more difficult for the methods to predict genes that are broadly expressed across all cell types/tissues. The genome context descriptors showed a weak positive association with model performance. Of all descriptors, the number of surrounding genes had the weakest influence, suggesting that genes in gene-dense regions are not more challenging than those without many other genes in their surrounding. A weak positive association was observed for the number of CREs (Fig. 3b), which was particularly pronounced for up to 300 CREs around a gene. Overall, we observed consistent effects of the gene descriptors across methods, with minor differences in association strength. There was also a dependency of the model performance on the biotype of a gene. Consistently across all models, the expression of protein-coding genes was predicted with a higher accuracy than of non-coding RNA genes or pseudogenes (Supp. Fig. 4b,c). Nonetheless, models trained for non-coding RNA genes and pseudogenes were still giving accurate predictions.

Additional insights were gained by investigating the difference in the distribution of gene descriptors of genes for which one method outperforms all others. For example, Fig. 3d shows the length distribution of the longest 5’ UTRs per gene where a method performed strictly better than the other methods. While we observed previously that method performance was positively associated with the 5’ UTR length, Binned-CNN performed particularly well when 5’ UTRs were long in comparison to other methods. The CRE-MLP method was best when many genes had zero expression (Fig. 3e) and many CREs (Fig. 3f), while both RF methods were best for genes with few test samples with zero expression. As RF approaches average over trees it is difficult to predict exactly zero. Overall, ML methods differed in their ability to extract useful information from H3K27ac data on gene expression.

### Model interpretation via *in silico* perturbation

The key to interpreting the models and revealing underlying regulatory mechanisms lies in the feature importance, meaning which regulatory regions are relevant to predict the expression of a target gene. As a measure of feature importance, an *in silico* perturbation (ISP) approach (Fig. 4a) was applied. For a target gene, each putative regulatory region was iteratively knocked out, i.e. each feature was set to zero, and the predicted expression value of the pre-trained model was obtained. The ISP was then calculated as log-ratio between wildtype and perturbed expression prediction. The ISP can be positive or negative, i.e. the ISP indicates not only the strength of the region-gene interaction but also whether the region has an enhancing or repressive effect on the expression of the target gene. The ISP was used as an alternative to other interpretation methods, such as SHAP [29], which require large computational resources and time, especially in our setup with a large number of models and features. Figure 4b displays the ISP tracks of the five approaches at the *LMO2* locus together with K562 H3K27ac ChIP-seq signal, experimentally validated CRE-*LMO2* interactions and chromHMM regions [28]. The different feature setups allow different levels of resolution. Overall, the methods largely agreed on the locations of putative and validated CREs, but not necessarily on the direction or strength of the regulatory effect. High ISP values overlapped with chromHMM states that correspond to active transcription and enhancers and were consistent with experimentally validated regulatory regions affecting *LMO2* expression in K562 cells [27].

**Fig. 4.**
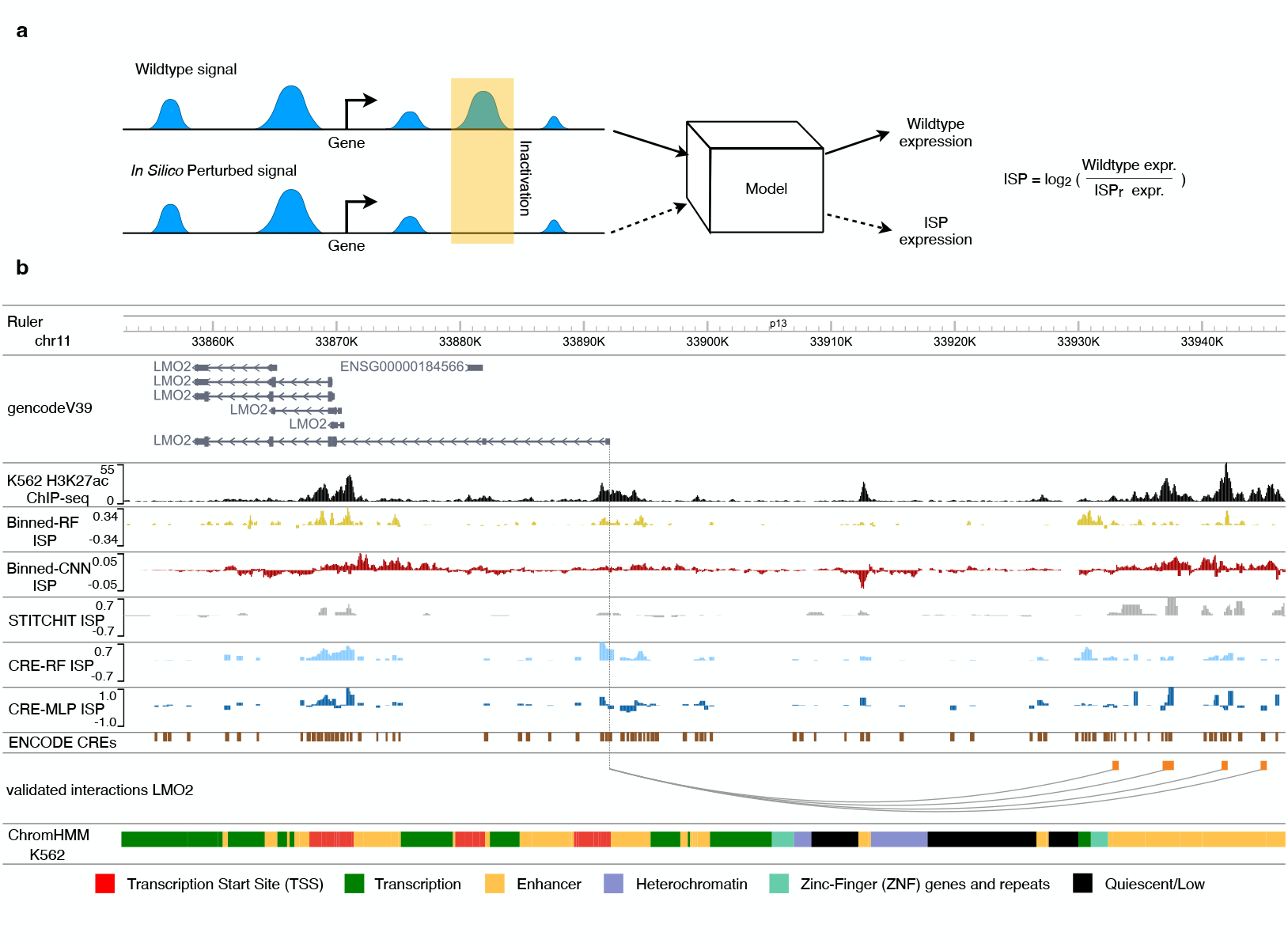
Illustration of the *in silico* perturbation approach. **(a)** *In silico* perturbation (ISP) compares the predicted gene expression using wildtype input signal (Wildtype expr), and expression predicted for input signal where one input feature *r* is set to zero (ISP_*r*_ expr). **(b)** WashU Browser [26] view of the K562 H3K27ac ChIP-seq signal track at the *LMO2* locus (*top*), followed by the K562 ISP tracks for each method, pre-defined candidate CREs from ENCODE [25], CRISPRi-validated interactions [27] and chromHMM states [28]. Note that the ISP scale is different for each method, thus not directly comparable. The EpiATLAS K562 sample was used as input data (IHECRE00001887). The chromHMM states TssA, TssFlnk, TssFlnkU, TssFlnkD were merged together into state *TSS*, Tx and TxWk into *Tx*, and EnhG1, EnhG2, EnhA1, EnhA2, EnhWk into *Enhancer*

### Model comparison on CRISPRi and eQTL data

The models were compared with regard to their ability to predict experimentally validated enhancer-gene interactions [27] by estimating the effect of an enhancer on a gene’s expression via ISP. The CRE-RF model achieved the highest performance with an area under the precision recall curve (AUPRC) of 0.38, followed by STITCHIT with 0.35 (Fig. 5a,b). The CRE-MLP model yielded the lowest AUPRC with a value of 0.33. For the models trained on the CRE features, normalizing the ISP score per gene increased the performance (Eq. 1 & 2, Supp. Fig. 5a). Overall, the variation in performance between the models was rather low, which suggests that the regression setup and the ISP approach are more defining for the detection of validated interactions than the specific feature and model setups.

**Fig. 5.**
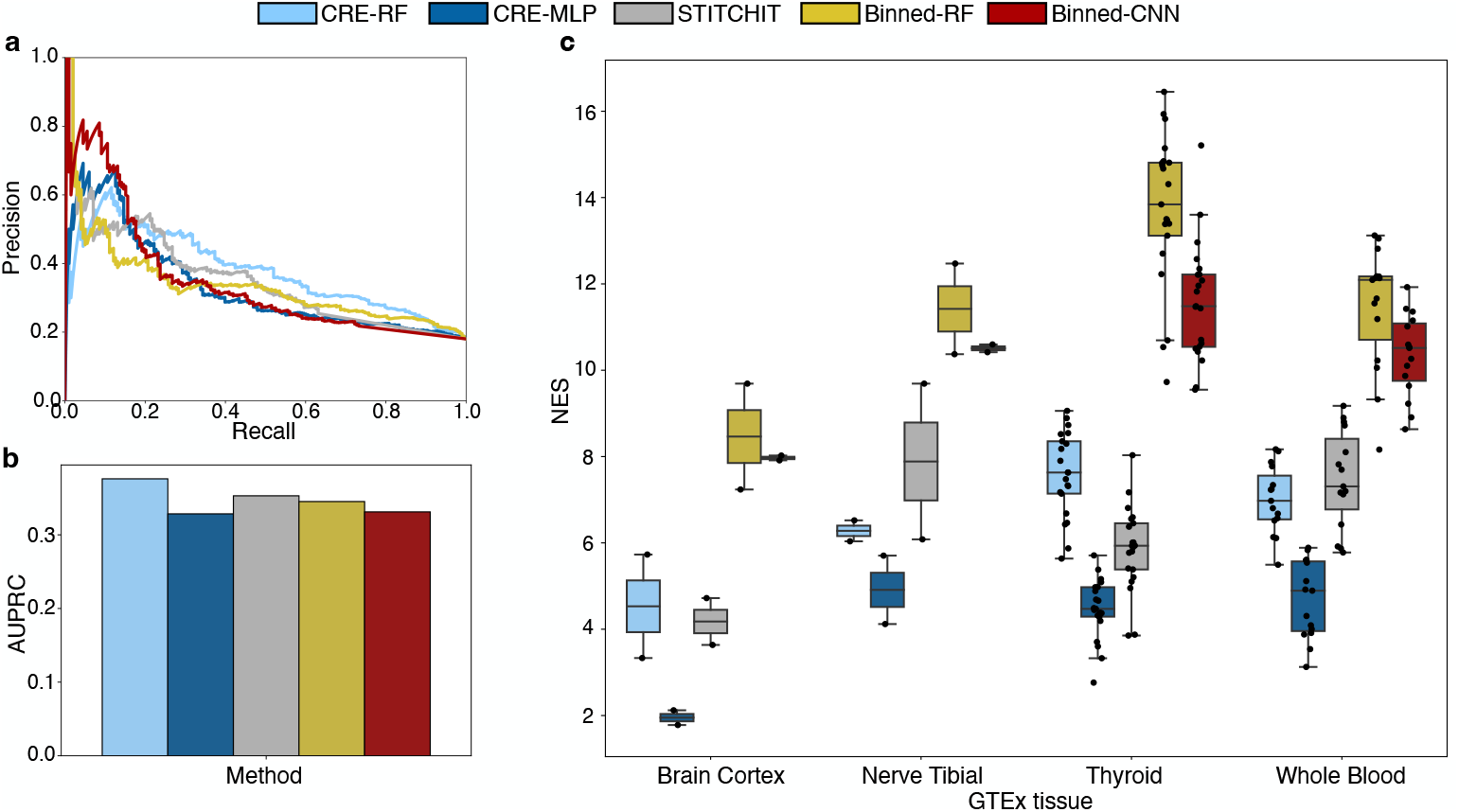
Method comparison using CRISPRi and eQTL data. **(a)** Precision recall curves of all models based on validated enhancer-gene interactions [27]. An *in silico* perturbation (ISP) approach was implemented to assess the importance of an enhancer-gene interaction for a model (illustrated in Fig. 4a). Two score calculations were tested: *ISP* and *ISP*_*normG*_ (Eq. 1&2). The better one is shown. The curves were created by using the set of interactions for which all models could produce a score (1,099 tested interactions out of which 198 were true positives). **(b)** Barplot with the area under the precision recall curve (AUPRC) for the curves in (a). **(c)** Normalized enrichment score (NES) of the models for enhancer-gene interactions that are supported by eQTL-gene pairs from GTEx [30–32]. Shown are the eQTL-gene pairs fine-mapped with DAP-G [33]. The boxplots are formed by the NES of the EpiATLAS samples that were matched to the respective GTEx tissue (center line median, boxlimits inter-quartile range, whiskers up to 1.5x inter-quartile range). Similar to (a), *ISP* and *ISP*_*normG*_ were tested and the one with higher average NES across all samples was kept. For other fine-mapping methods and score calculations see Supp. Fig. 5.

Another source for validation were expression-quantitative trait loci (eQTL) gene interactions. When compared on eQTL data, the models with the binned feature setups had the highest enrichment of eQTL-supported enhancer-gene interactions among their top ranked interactions (Fig. 5c). Binned-RF achieved the highest enrichment for 82.5% of all tested samples. Among the CRE-models, CRE-RF had higher enrichment scores than CRE-MLP. For all models the enrichment was higher when the score was not normalized per gene. The results for the three tested eQTL fine-mapping methods were highly similar (Supp. Fig. 5b-d).

### A novel concept for conducting histone-acetylome-wide association studies

The availability of accurate prediction models enabled a novel application for conducting a histone-acetylome-wide association study (HAWAS) with H3K27ac data for the discovery of disease genes. We suggest a new model-based approach to first find genes associated with the disease and then the relevant H3K27ac signal regions (Fig. 1d). First, a trained ML model is used to predict the expression in control and disease samples for over 28,000 human genes. Then, an association test per gene is performed, to identify genes that show significant alteration in their predicted expression (HAWAS-gene). Second, for these candidate disease genes, the ML model is used to estimate the importance for all gene features by calculating the ISP values. These are used for a second association test to identify the regulatory regions (features) of the ML model where the ISP values differ between control and disease samples (HAWAS-region).

The trained models from the two best-performing methods (CRE-RF, Binned-CNN) were applied to Chronic Lymphocytic Leukemia (CLL) patient data. CLL is known to lead to large differences in epigenomes and transcriptomes of patient cells [37, 38]. After conducting the HAWAS-gene test, different numbers of significant genes were identified for each approach: 13,473 for Binned-CNN and 16,626 for CRE-RF. For comparison, differential analysis of gene expression with measured RNA-seq data from the same biological samples resulted in 14,998 genes and large overlaps with the computational methods (Fig. 6a). This suggests that both ML approaches captured relevant biological differences in expression. Strikingly, the CRE-RF based analysis identified 97% of the genes obtained from RNA-seq analysis. Indeed, the predicted fold changes between healthy and control samples were most similar for the CRE-RF (Fig. 6b, R=0.96) and less accurate for the Binned-CNN (Fig. 6c, R=0.90). Enrichment analysis with known CLL genes (DisGeNET database [34]) indicated a significant overlap for each approach (Fig. 6d), suggesting that the models were successfully identifying genes involved in CLL pathogenesis.

**Fig. 6.**
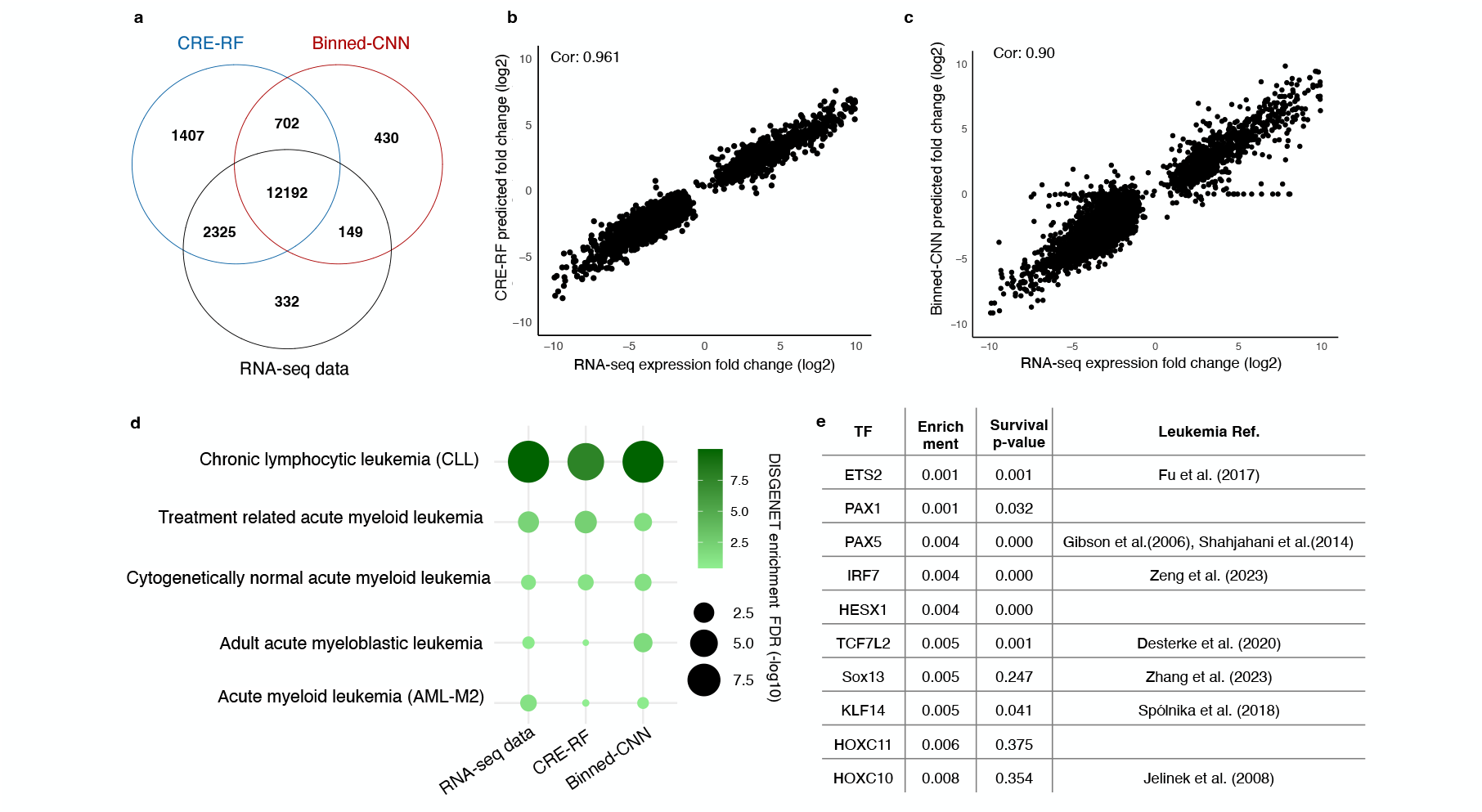
Model-based histone-acetylome-wide association study using pre-trained ML methods reveals insights into leukemia gene regulation. **(a)** Venn diagram showing the number of significantly associated CLL genes for Binned-CNN, CRE-RF, and experimentally measured RNA-seq data from the same individuals. **(b**,**c)** Scatterplot of gene expression fold change values (log2) for real RNA-seq measurements versus predicted for 14,998 genes (differentially expressed in real data) (Pearson correlation indicated in the plot). Comparisons shown for CRE-RF **(b)** and Binned-CNN **(c)** predictions. **(d)** Dot plot illustrating the overlap of known leukemia genes (Dis-GeNET database [34]) and CLL genes detected by CRE-RF, Binned-CNN, and RNA-seq expression analysis (enrichment test -log10 FDR). **(e)** Table showing the top 10 enriched transcription factors (TFs) identified by motif enrichment with PASTAA [35] on 49,920 CREs associated with CLL using CRE-RF models. PASTAA enrichment FDR and supporting references are shown in the second and last columns. Survival p-value is obtained from Kaplan-Meier survival analysis with log-rank test [36].

We also compared the HAWAS gene-test to conventional approaches for associating histone modification changes to genes. We took regions with differential signal (n = 223,586) and associated them either to the nearest gene (n = 22,409) or to their putative target genes according to regulatory interactions predicted with the generalized Activity-by-Contact (gABC) score (n = 25,145) [39, 40]. The conventional approaches were less specific and found thousands of genes without a significant expression change. For both conventional approaches, 56% of the identified genes were differential according to the RNA-seq data, while 87% of the genes found by CRE-RF and 91% of Binned-CNN genes were differentially expressed. As a consequence, genes found by the HAWAS-gene tests had stronger expression changes (Supp. Fig. 7).

Because CRE-RF was the most accurate in predicting differential RNA expression, it was used for the HAWAS-region analysis to uncover 49,920 significant regions from the 16,626 significant genes (*FDR* ≤ 0.05). 45.1% of the identified regions overlapped sites that were significant in a direct comparison of the ChIP-seq signal between control and disease [39]. Significant variation in active euchromatin may affect the binding of transcription factors (TFs) in those regulatory regions. A TF motif enrichment analysis [35] of the CRE-RF identified regions revealed 30 associated TFs (FDR 0.01). Among the top 10 TFs, 7 were found to be relevant to leukemia according to literature evidence (Fig. 6e) [41–48]. Kaplan-Meier survival analysis with log-rank tests (median cut point) was conducted for the top 10 TFs between control and CLL samples from established cohorts using the Survival Genie2 tool [36]. Seven out of the ten TFs had a significant survival difference with *p*-value ≤ 0.05 (Fig. 6e and Supp. Fig. 8). Thus, performing the HAWAS on the model predictions provided valuable insights into the underlying genes and regulating TFs in CLL, highlighting the utility of the developed models for biomedical research applications.

## Discussion

The EpiATLAS resource was used to train and benchmark various machine learning approaches in a gene-specific learning setup. A challenge for the comparison was that a method required individual optimization for over 28,000 genes. Naturally, there was a limitation in the variety of model topologies and parameter setups that could be explored in this study. We realized that the feature setup played a crucial role for model performance. The advantage of CRE-RF, with predefined regions, over the Binned-RF could originate from the removal of many irrelevant features before learning. Using regularization may be an alternative way to improve performance of the RF-based approaches [49]. Further, there are many additional topologies and components for neural networks that one could try. We expect that using the EpiATLAS dataset further improvements will be made in the future. Also, it is likely that improvements can be made by adjusting for gene descriptors that affect a model’s performance (Fig. 3).

It is reasonable to assume that expression prediction for a gene that shows a complex regulatory pattern (many CREs, nonlinear combinations) is difficult at the current size of 792 training samples. To improve convergence for CNN models, we used a warm-start initialization procedure, which increased accuracy for many genes, but applying a similar strategy was not feasible for MLP models, as their feature space differed between genes. Exploration of other ideas-such as adjusting model complexity, separating genes with low or high average expression, or testing other deep learning architectures-are directions for future work.

Methods from explainable AI [29, 50] are commonly used to investigate the contribution of individual features for prediction outcome. Since over 28,000 models were investigated, the computationally efficient ISP approach was used to assess the impact of a genomic region on the expression prediction of a particular model. The CRE-based models use predefined genomic regions, while STITCHIT and the binned models start with the features from the entire window. Consequently, only the latter models can find regions of interest that are outside of predefined CREs. New model-based applications were suggested in the context of HAWAS that estimate genes or regulatory regions of genes associated with a disease. In particular, the approach for model-based transcriptome-wide association studies (TWAS) [24, 51] is conceptually similar, except that in a TWAS disease genes are predicted from genomic variant data, not from epigenome data.

Applying the model-based HAWAS to data from leukemia patients revealed good agreement with RNA-seq differential gene expression analysis and identified regulators associated to survival, even at a relatively small cohort size of 22 samples (Fig. 6). Thus, for future HAWAS the experimental efforts could be limited to measuring only the epigenome, which may be important in situations where measuring RNA transcripts is complicated or may allow lowering sequencing costs. However, it can be expected that the application of the models on cell types that were not part of the EpiATLAS will show reduced performance, particularly for cell type-specific genes. Currently, our models are available for over 28,000 human genes and larger data sets are needed to learn models for genes that had too little variation in the training set. The ability to obtain disease genes and their CREs using case-control epigenome data as illustrated here, will fuel future association studies and reveal cell type-specific roles of the epigenome in disease.

## Methods

### EpiATLAS epigenome and transcriptome data

Model training was based on the data of the International Human Epigenome Consortium (IHEC), available via the EpiATLAS portal (https://ihec-epigenomes.org/epiatlas/data/) [23]. The activity of genomic regions was assessed by the H3K27ac ChIP-seq signal, more precisely the average fold-change signal over the background in a given region. For gene expression the expected counts from RSEM [52] were used after size factor normalization with DESeq2 [53]. All 965 samples were included that had both H3K27ac ChIP-seq and RNA-seq available, according to the metadata version 1.1. As cell type/tissue label for the samples, the column *harmonized_sample_ontology_intermediate* was used, which encompasses 51 different ontology terms for the 965 samples. If multiple RNA-seq experiments were provided per sample, the total RNA-seq experiment was chosen. All data is in human genome version GRCh38. As gene annotation we used v38 from GENCODE [54].

### Dataset filtering and preparation

We filtered for genes with ≥ 2 expression variance and 90% non-zero values across samples in the RNA data. This resulted in a set of 28,180 genes that we retained for the analyses throughout this study. Supp. Fig. 4b provides details on the biotypes of the retained genes. For all methods, the feature and response matrix were *log*2-transformed with a pseudo-count of 1. If additional scaling was performed, it was first applied to the training data, and the scaling factors from the train data then utilized to normalize the test data to prevent information leakage. Details on the scaling procedure used by each method can be found in the corresponding sections.

### Feature setups

To define the candidate regulatory regions for a gene, three distinct strategies were employed: *CRE, binned*, and *STITCHIT*. For the *CRE* strategy, candidate CREs from ENCODE were used to identify putative regulatory regions within the 1 MB window surrounding the TSS. More precisely, the candidate enhancers and candidate promoters were downloaded from ENCODE’s SCREEN web interface (v3) [25] and merged into a joint set of 988,415 candidate CREs. In the *binned* approach, the abundance of H3K27ac read counts was calculated in consecutive 100 bp bins spanning a 1 MB window symmetrically centered on the 5’ TSS of the gene. Similar to the binned approach, for *STITCHIT* ‘s feature selection the genome was divided into equal-sized bins holding the mean H3K27ac signal in a 1 MB window around the TSS of a target gene. The bins of size 10 bp were successively merged into larger segments within the STITCHIT algorithm. Figure 1b schematically depicts the three feature designs utilized in this study.

### Data partitioning and cross validation for training

To benchmark the performance of the different approaches, the dataset was randomly partitioned into a training (80%) and test set (20%). All reported model performances were calculated on the test partition. For all methods except RF, a cross-validation (CV) procedure was applied for parameter-optimization on the training data. The best model was selected according to the minimum CV-error. The CV-error corresponds to the average mean-squared error (MSE) over the CV-folds. Details on the CV-parameters can be found in the respective method section.

### Setup for Machine Learning models

#### CRE-RF

Random Forest model was trained using the randomForest function from randomForest library version (4.7.1.1) in R. The data was normalized to the range [0,1] using min-max scaling. The analysis was carried out using the following arguments without any hyperparameter tuning:

~~~
  ntree = 501, mtry = floor(sqrt(ncol(training))),
  replace = TRUE, nodesize = 5,
  maxnodes = NULL, importance = TRUE,
  localImp = FALSE, proximity = FALSE,
  oob.prox = FALSE, norm.votes = TRUE,
  do.trace = FALSE, keep.inbag = FALSE
~~~

#### CRE-MLP

Two different designs were explored for the MLP models: one with a single hidden layer and another with two hidden layers (Supp. Fig. 2a,b). The single hidden layer MLP consisted of 100 nodes, while the two hidden layer MLP comprised 100 nodes in the first layer and 50 nodes in the second layer. ReLU activation functions were used for all hidden layers, while a linear activation function was applied to the output layer. Hyperparameter tuning was conducted for each topology to identify the most effective model configurations. Given the high computational cost of exhaustive hyperparameter search, parameter combinations were evaluated on a subset of 500 genes. A search grid for initial seed (30 random seeds), dropout rates (0.2 and 0.4 for both the first and second hidden layers), batch size (fixed at 32), maximum epochs (set to 1200 with an early stopping criterion of 30 epochs patience), and default learning rates for Adam optimizer (at 0.001) was defined [55]. The data was normalized to the range [0,1] using min-max scaling, then MLP models were trained using the *Keras* library in R with the help of its built-in validation features. The validation set accounted for 30% of the training data and was used to assess model performance during training.

#### Binned-RF

We used the Random Forest for regression on the binned data by utilizing the `RandomForestRegressor` function from the scikit-learn library in Python. The response was min-max scaled to the range [0,1]. The analysis was carried out using the following arguments:

~~~
n_estimators=100, criterion=‘squared_error’,
max_depth=None, min_samples_split=2,
min_samples_leaf=1, min_weight_fraction_leaf=0.0,
max_features=1.0, max_leaf_nodes=None,
min_impurity_decrease=0.0, bootstrap=True,
oob_score=False, n_jobs=None,
random_state=0, verbose=0,
warm_start=False, ccp_alpha=0.0,
max_samples=None, monotonic_cst=None
~~~

#### Binned-CNN

Given the nature of the features in the binned setup, a convolutional neural network (CNN) was selected. Two primary CNN topologies were employed in this study: the contracting kernel and the expanding kernel (Supp. Fig. 2c,d). In the contracting kernel, the kernel sizes gradually decrease from the input layer to the output layer. Conversely, the expanding kernel topology features progressively larger kernel sizes. For both topologies, a dense layer with 5 hidden nodes connects the convolutional block to the output regression node. For all models, we tested the CNN results by varying the activation functions in the first and last layers. Specifically, we explored the influence of ReLU and sigmoid activation functions in these layers. Since the initialization of weights was crucial for optimization, a heuristic warm start strategy was developed to initialize the weights. Inspired by the concept of transfer learning, three distinct gene pools were created, each serving as a separate dataset. Specifically, batches of 1,000 genes were randomly selected from the entire gene set, along with their corresponding biological samples, to form one of three datasets. The contracting and expanding kernel topologies were trained on these three datasets using the ReLU activation function throughout. Each gave an optimal model for the 1,000 gene set and the corresponding parameters of each of these three models were used as initial weights (warm start) to train the CNN models in a gene-specific manner. For genes located near the chromosome borders, where a complete 10,000-bin setup was not feasible, we could not transfer the parameter initializations and we thus trained the CNN models using a cold start. Before model training, the response was min-max scaled to the range [0,1]. The features were scaled to unit-variance using a batch normalization layer. All CNN implementations were carried out using the *Keras* library in Python, with a validation split of 0.2 and a learning rate of 0.001. The models were trained using the Adam optimizer for up to 1000 epochs [55], with batch size of 32 and early stopping applied to prevent overfitting (patience parameter set to 5). For both contracting and expanding CNN models, the stride in the first convolutional layer was set to 5. In the expanding model, subsequent layers used a stride of 2, while in the contracting model, a stride of 1 was applied to all remaining convolutional layers. All other parameters were kept at their default settings.

### STITCHIT

STITCHIT incorporates the identification of putative regulatory elements in a segmentation step before applying model learning. The initial bins were merged into larger segments such that the H3K27ac signal variance within a segment is low across all samples, with respect to discrete gene expression labels. The gene expression labels comprised binary classes {0, 1} that represent `expression` and no `expression` generated by a sample-wise discretization approach described in [56]. The segments that were associated with changes in the continuous expression of the target gene were chosen for the subsequent linear learning step. For this purpose, the Pearson correlation was calculated between the H3K27ac signal and the expression of the target gene across all samples. All segments that passed a significance threshold of *P* ≤ 0.05 were selected as features for predicting gene expression using an elastic net model. A six-fold nested CV-procedure was employed for parameter tuning and model selection on the standardized training data. Details of the STITCHIT algorithm are available in [22].

#### *In silico* perturbation score calculation

As an estimate for how important a genomic region is for the gene expression prediction in a sample, we calculated an *in silico* perturbation score (ISP). To do so, the signal of a given region *r* was set to zero in the input matrix and the expression predicted with this perturbed input. The ISP score is then defined as the *log*_2_-ratio of the predicted expression on the unperturbed data (*Wildtype expr*) over the perturbed expression prediction (*ISP*_*r*_*expr*):

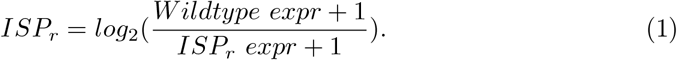

Since the models use different genomic regions as features, the regions that are perturbed for the ISP also differ. For the CRE-based models the individual CREs were set to zero and for STITCHIT the putative regulatory elements that it defines per gene were perturbed. Due to the large feature space of the binned models, multiple adjacent bins were perturbed together. More precisely, for a genomic region of interest, bins overlapping this region were set to zero, as well as the neighboring bins for a minimum of 10 bins. To summarize, the score can be calculated for each input feature of each gene model and each sample.

In addition, a score normalized per gene was tested for each model, which was calculated as follows:

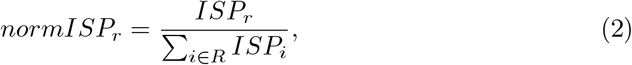

where *R* denotes all features of the gene. Thus, *normISPr* sets the effect that the removal of *r* has on the gene expression prediction in relation to the effect of removing any other feature of the gene. For models with a binned feature setup, this approach was not feasible due to the large feature space. This is why for the binned models, all candidate enhancers derived via thresholding their feature importance were taken as *R* (noted as feature fusion).

#### Feature fusion

Given that the global feature set for the Binned-CNN models spans the entire 1 MB genomic region divided into consecutive bins, an ad hoc procedure was implemented to derive ISP for these models. This approach included the following steps:

1. **Feature Importance Derivation:** Random Forest (RF) models were used to determine global feature importance on a per-gene basis.
2. **Binarization via Otsu’s Thresholding:** Feature importance values were binarized using Otsu’s thresholding method, categorizing bins as “active” (1) or “inactive” (0). The *threshold_otsu* function from Python’s *skimage*.*filters* package was used to calculate Otsu’s threshold.
3. **Fusion of Adjacent Bins:** Consecutive active bins were fused if separated by no more than five inactive bins.
4. **Expansion Strategy:** To ensure adequate impact on model prediction following ISP, fused regions shorter than 1 kb were expanded to a 1 kb span. This expansion sometimes resulted in merging adjacent fused regions, creating larger contiguous regions.
5. **Signal Nullification in Fused Regions:** For each expanded region, the signal in corresponding bins was nullified to calculate the effect of the perturbation on predicted gene expression.
6. **Prediction Ratio Calculation:** The ratio of the original prediction to the ISP prediction was calculated as per the formula outlined in Eq. (2).

Supplementary Fig.6b provides a visual summary of this procedure.

### Comparison on CRISPRi data via *in silico* perturbation

As experimentally validated enhancer-gene interactions, the data of Gschwind et al. was used, which is a reanalyzed collection of three CRISPRi-screens in K562 cells [57–59]. More precisely, the file EPCrisprBenchmark_ensemble_data_GRCh38.tsv.gz from their GitHub repository was taken, and all interactions that were labeled as significant in the *Significant* column of the table were considered true interactions. Repressive interactions, where the gene expression increased after enhancer perturbation, were not excluded. Additionally, the interactions were filtered for those with a distance between enhancer and the gene’s most 5’ TSS of ≤ 500 kb, matching the gene window in our setup. Mapping gene names to Ensembl IDs was done with the API of MyGene.info [60–62]. The resulting set of enhancer-gene interactions is available as supplementary material JointValidatedInteractions_hg38_500kb.txt on Zenodo. Processing of genomic coordinates was done with pybedtools (v0.9.0) [63, 64].

Since all CRISPRi data come from K562 cells, the ISP was done in the EpiATLAS K562 sample (IHECRE00001887). The ISP score was calculated for the genomic regions that are used as input for the models that overlap the validated CRISPRi regions (Eq. 1). Due to the different feature setups, the different models can produce ISP scores for different subsets of validated interactions. The models with a binned feature setup can score all validated interactions, while the CRE-based models and STITCHIT can only generate scores for interactions that overlap their features. The ISP score that is normalized per gene (Eq. 2) was tested as well.

All ISP scores were converted to their absolute values, since the validated interactions contain repressive and activating interactions. To construct PR curves for each model, a score cutoff was increased from the lowest to the highest absolute score in 10,000 steps. All interactions with an absolute score above the cutoff were considered as predicted positive interactions, and all below the cutoff as predicted negatives. Precision and recall were calculated for each cutoff based on the CRISPRi-validated interactions. If multiple features overlapped one CRISPRi-perturbed region, the absolute score of the features was summed.

### Ranking of eQTL-gene pairs

To compare how well the models recover interactions supported by eQTL-gene pairs, tissues from GTEx were mapped to matching samples from the EpiATLAS. Four tissues from GTEx were selected, two with a high number of matching IHEC EpiATLAS samples and two with only two matching samples (Tab. 1). Data of the three fine-mapping methods available on GTEx (dbGaP accession number phs000424.v8.p2) [32]) were used by accessing the following files: ‘CAVIAR_Results_v8_GTEx_LD_HighConfidentVariants.gz’ for CAVIAR [65], ‘GTEx_v8_finemapping_CaVEMaN.txt.gz’ for CaVEMaN [66] and ‘GTEx_v8_finemapping_DAPG.CS95.txt.gz’ for DAP-G [33]. For each of the models and each of the EpiATLAS samples, a normalized enrichment score was calculated to assess whether interactions supported by eQTL data were ranked higher than those not supported by eQTL data. This was repeated for each fine-mapping method. Each possible enhancer-gene pair for all genes with at least one eQTL in a tissue was scored, meaning each input feature of a model was iteratively perturbed to calculate the ISP for a given sample (Eq. 1 and Eq. 2). For the models with the binned feature setup, the candidate enhancers were defined by thresholding the feature importance of the bins. The absolute scores were subsequently sorted in descending order. To consider the same number of interactions across all methods, the 100,000 highest ranked interactions were taken for each EpiATLAS sample and each model. To also have the same number of interactions supported by eQTL-gene pairs for each method, the set of overlapping eQTL-gene pairs were reduced to the same size per sample (minimum of overlap across methods). Based on the 100,000 interactions, a normalized enrichment score was calculated with the prerank function of GSEAPY (v.1.1.2) [67], which is a Python implementation of GSEA [30, 31]. The normalized enrichment score is the ratio of the enrichment score and the average enrichment score across random dataset permutations. The number of permutations was set to 100. The weighting parameter p was set to zero, so that the calculated enrichment score equals a Kolmogorov-Smirnov statistic.

**Table 1.** Overview of matched IHEC samples for each of the GTEx eQTL sets. GTEx tissues are matched to EpiATLAS biosamples according to the EpiATLAS metadata file (v1.1). For a mapping of the individual samples see supplementary material IHEC_ValidationSamples.txt.

| GTEx tissue | EpiATLAS biosample | # samples | EpiATLAS metadata column used |
| --- | --- | --- | --- |
| Brain_Cortex | brain | 2 | harmonized_tissue_type |
| Nerve_Tibial | tibial nerve | 2 | harmonized_tissue_type |
| Thyroid | thyroid gland | 21 | harmonized_tissue_type |
| Whole_Blood | venous blood | 15 | harmonized_sample_ontology_intermediate |

### Partial correlation analysis

To assess the relationship between gene-specific attributes and the models performance, a partial correlation analysis was conducted across multivariate input arguments using the psych package [68], specifically employing the partial.r function in R. In this analysis, descriptors were organized into three categories: The *structural* descriptors correspond to: i) the number of TSSs, ii) the gene length including introns (gene length), iii) gene length excluding introns (exon length) and the lengths of the longest iv) 5’ and v) 3’ UTRs. The *expression* group comprises i) the proportion of samples in the test set in which the expression of the target gene is nonzero and ii) the expression breadth.

For the expression breadth, all RNA-seq samples from the IHEC EpiATLAS (including those without matching H3K27ac CHiP-seq data) were used and the expression averaged across all samples belonging to a cell type/tissue (|*C* | = 58). To then get an estimate of how broadly a gene is expressed, the fraction of cell types/tissues in which the average expression *µ*_*g,c*_ exceeds 0.5 TPM was taken:

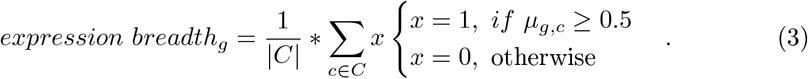

The *genome context* descriptor set holds: i) the number of ENCODE CREs and ii) the number of genes excluding the target gene. Both descriptors were counted within a 1 MB window around the target gene. All descriptors were obtained from the GENCODE v38 annotation file [54].

The partial correlation values between model performance and each descriptor were calculated while conditioning on the other descriptor categories in order to compare the contribution of each descriptor within its respective group (Supp. Fig. 4a). For example, for number of TSSs the correlation was calculated as follows:

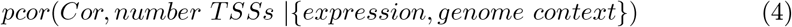

where Cor denotes the Pearson correlation of the model on test data and {*expression, genome context*} subsumes all descriptors in these two descriptor categories.

In this approach, conditioning variables included only genomic descriptor of other categories. This process was repeated iteratively to obtain rank-based Spearman partial correlation values for each descriptor from each of the three categories, allowing us to assess their unique contributions to model performance.

### HAWAS-gene test

The HAWAS-gene approach leverages ML-based predictions of gene expression to identify genes with significant alterations between healthy and disease states. By comparing the predicted expression profiles of thousands of genes, this approach enables the discovery of potential disease-associated genes driven by epigenetic regulation (Algorithm 1).

In the first step, two best pre-trained ML models (CRE-RF and Binned-CNN) are used to predict gene expression counts based on H3K27ac signal data from both control and disease samples. For *z* genes, there are *z* corresponding models, where each model *M*_*i*_ (*i* ∈ {1, 2, …, *z*}) is associated with an input matrix *G*^*n×m*^. Here, *n* represents the total number of samples, including both control and disease groups, and *m* represents the number of genomic features (regions) of the gene model *Mi*.

For each gene *i*, the model produces the predicted expression count and stores it in *E*_*s,i*_. This process is repeated for all *z* genes, and the final results are stored in the output matrix *E*^*n×z*^, where *n* (rows) corresponds to the input samples, and *z* (columns) represents the predicted expression values for all genes.

In the second step, differentially expressed genes between control and disease samples are identified using DESeq2 [53]. Specifically, a design matrix is constructed to model the two conditions: control and disease. DESeq2 is then applied to the predicted expression count matrix *E*^*n×z*^. The test returns a list of disease-associated genes based on statistically significant expression changes.

#### Algorithm 1

HAWAS-gene test

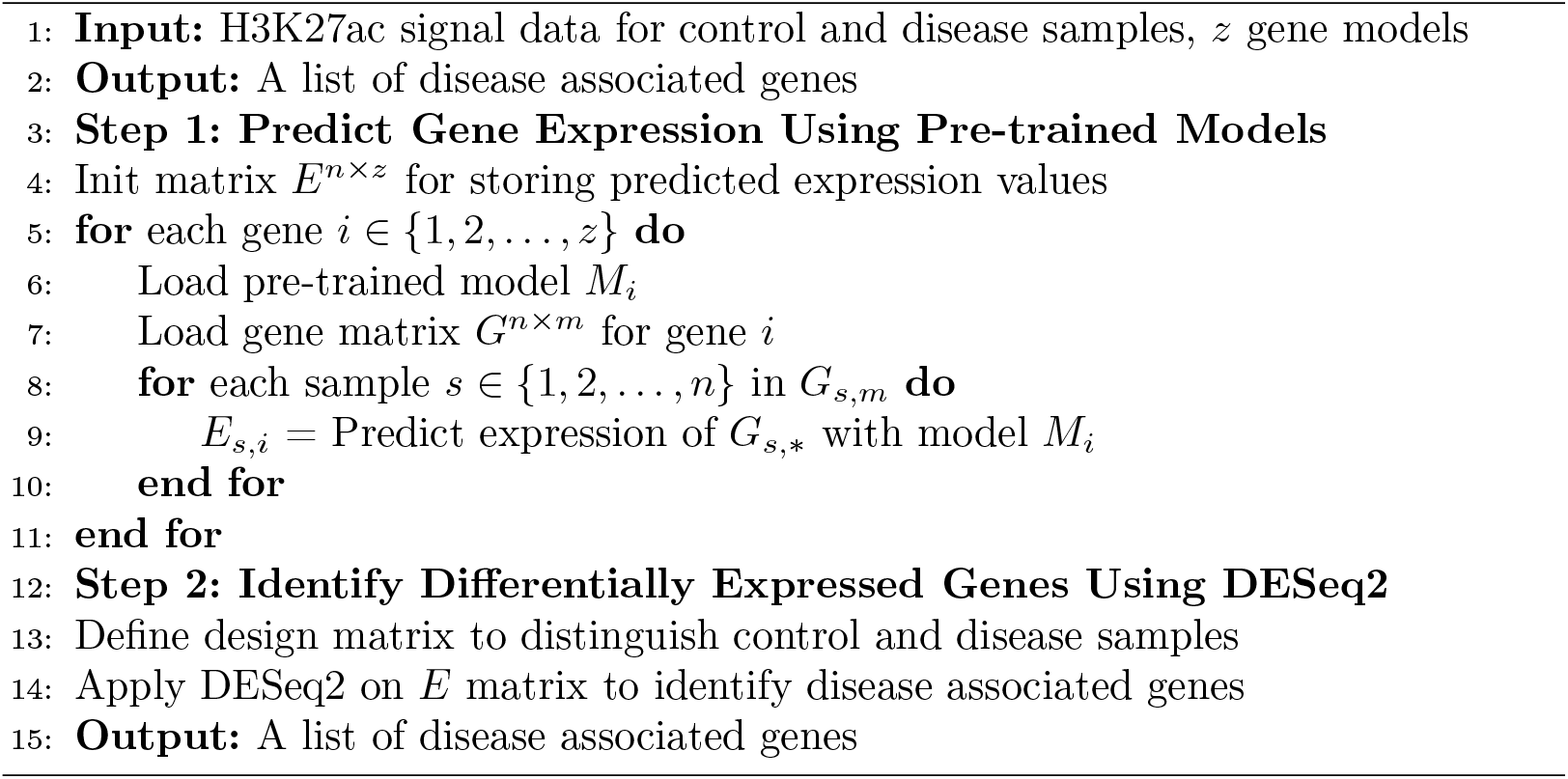

### HAWAS-region test

While identifying HAWAS genes provides critical insights, understanding which regulatory elements contribute to these expression changes is essential for deciphering disease mechanisms. The HAWAS-region test extends the analysis to the level of gene regulatory regions by quantifying the contribution of individual regions (features) to gene expression by using ISP. This method identifies regulatory elements whose influence on gene expression differs between healthy and disease conditions (Algorithm 2).

In detail, the input of this method are *z* gene models. The input for each gene *i* is a *G*^*n×m*^ matrix with H3K27ac signal with *n* rows representing samples and *m* columns representing regions. In the first step, the ISP score is calculated for each sample *s* ∈ {1, 2, …, *n*} for all regions *r* ∈ {1, 2, …, *m*} of gene *i* using the ISP formula (Eq. 1) resulting in a matrix *I*^*s×r*^. For notational simplicity we assume here that each gene *i* has the same number of regions *m*. In the second step, a *t*-test is performed between the control and disease groups for gene *i* across all *m* regulatory regions and the resulting *p*-values are stored in *L*^*i×m*^. *t*-test is chosen because it is well-suited for comparing continuous values, e.g., region importance scores. A two-sided test is used. Hence, after the *t*-test there are *m* many *p*-values for the gene *i*. This process is repeated for all *z* genes and there exist *z* × *m* many *p*-values that are stored in list *L*. In the third step, a false discovery rate (FDR) correction is applied on the *p*-values from list *L* to adjust for multiple testing [69]. Hence, the output is list *W* which contains significant regions with their adjusted *p*-values.

### Transcription factor enrichment and survival analysis

To investigate the potential regulatory mechanisms underlying disease-associated regions, TF motif enrichment analysis was performed using the PASTAA algorithm [35]. This analysis was applied to the significant regulatory regions identified by the HAWAS-region test using the CRE-RF model. Enrichment results were filtered based on a false discovery rate (FDR) threshold of 0.01, and top-ranking TFs were further examined based on literature evidence for relevance to leukemia.

For survival analysis, the Survival Genie2 tool [36] was used, and lymphoid leukemia was selected as the disease of interest. We chose the TARGET-ALL-P2-Bone-Marrow dataset, with overall survival as outcome type, took primary tumor types and considered all available samples. The top 10 TFs identified from the motif enrichment analysis were evaluated across these datasets. Patients were stratified based on the median expression levels of each TF, and survival differences between groups were assessed using the log-rank test. TFs with *p*-values ≤ 0.05 were considered significant.

### HAWAS on CLL data

Twenty-six adult samples, consisting of 12 healthy and 14 CLL samples (including both female and male), were initially considered for the HAWAS-gene test to determine CLL-related genes. However, three samples were removed as outliers based on PCA analysis, and one sample was excluded due to unknown sex. After filtering, 22 samples (10 healthy and 12 CLL) remained for analysis. There were some modification in the pipeline based on the data characteristics. First, for each of the 27,385 shared genes, CRE-RF and Binned-CNN models were used to predict expression values in the selected samples. Second, before performing the DESeq2 analysis, genes with fewer than 10 counts across all samples were removed. Third, since the size factor-normalized read counts are used in this study the size factor variable in the DESeq2 pipeline was set to 1 to prevent normalization. Fourth, sex was included as a covariate in the analysis. The metadata predicted by EpiClass was used for the covariate, more precisely the column “EpiClass_pred_Sex”.

#### Algorithm 2

HAWAS-region test

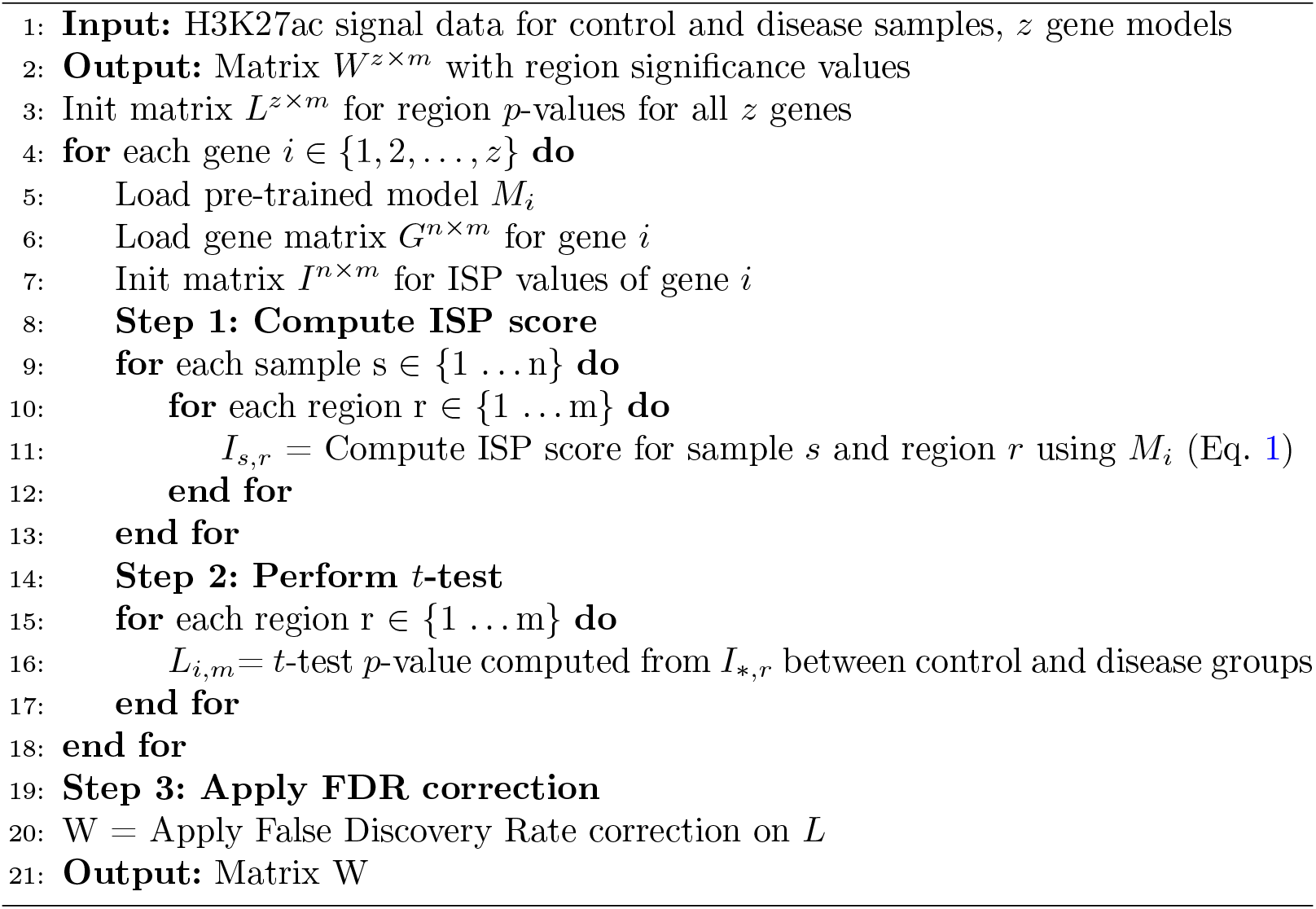

To determine related regions (features) of the significant genes of the HAWAS-gene test, the HAWAS-region test was used with an adjusted *p*-value cutoff of 0.05. In cases where redundant regions remained after FDR correction, the one with the smallest adjusted *p*-value was selected. The found HAWAS regions were further analyzed to uncover potential TFs involved in disease-specific gene regulation. For this, PASTAA [35] was used with an FDR threshold of 0.01. As TF motifs, a collection of 818 position frequency matrices from JASPAR 2022 [70], HOCOMOCO v11 [71] and the work of Kheradpour and Kellis [72] were used, available under https://github.com/SchulzLab/STARE/blob/main/PWMs/2.2/Jaspar_Hocomoco_Kellis_human_transfac.txt. Moreover, to assess the biological relevance of the HAWAS genes, a comparison was made to known CLL genes from the DisGeNET database [34] (Fig. 6d).

### CLL-genes via nearest gene and gABC

To have a comparison of the HAWAS-gene test to conventional approaches, regions with differential H3K27ac signal between control and CLL were called with DiffBind (v.3.4.11) [39]. As peaks, the ENCODE CREs were taken and the ChIP-seq input was given as background for each sample. Peaks with FDR ≤ 0.05 and an absolute log fold-change of ≥ 0.585 were considered differential. To then link the differential peaks to genes, two approaches were used. First, each differential peak was associated to its nearest gene within 100 kb based on the distance to the 5’ TSS with the closest function from bedtools [64]. Second, differential peaks were mapped to all genes they form interactions with according to the gABC-score. To get the gABC-score, STARE (v.1.0.4) [40] was run for each condition with all ENCODE CREs as candidate enhancers and their average fold-change H3K27ac ChIP-seq signal as enhancer activity. As contact data, an average Hi-C matrix was used [27]. The window size was set to 1 MB, the score cutoff to 0.02 and regions known to accumulate anomalous signal were removed [73, 74]. If a gene was predicted to form an interaction with a differential peak in either condition, it was added to the gABC-score gene set. For both approaches, only genes that were also considered for model learning were kept.

## Supplementary information

A supplementary file containing additional figures is available. These figures provide further details and support the results presented in the main text of the article.

## Author contributions

D.H. collected and prepared the IHEC data and implemented and performed the validation on CRISPRi and eQTL data, and comparison to conventional HAWAS approaches. S.A. implemented and ran the CRE-RF and CRE-MLP models and implemented and performed the HAWAS analyses (genes test, regions test, TF enrichment and survival analysis, HAWAS on CLL data). L.R. adjusted and ran STITCHIT. F.B.A. implemented and ran the Binned-RF and Binned-CNN models. The project was designed and supervised by M.H.S. Manuscript writing was done by all authors.

## Data availability

The trained models of the two best performing approaches (CRE-RF and Binned-CNN) are available on Zenodo at the following link: https://zenodo.org/uploads/13992024. Training data was taken from the IHEC EpiATLAS portal (https://ihec-epigenomes.org/epiatlas/data/). All other data was taken from public resources as indicated in the methods.

## Code availability

The code used for the analyses is publicly available on GitHub at the following repository: https://github.com/SchulzLab/EpiExpress. It includes scripts to use pre-trained models on custom input data.

## Acknowledgements

We would like to thank Jonas Fischer and Ivan Costa for their valuable feedback on the manuscript.

