## Supplemental_figures for "Harnessing machine learning models for epigenome to transcriptome association studies"

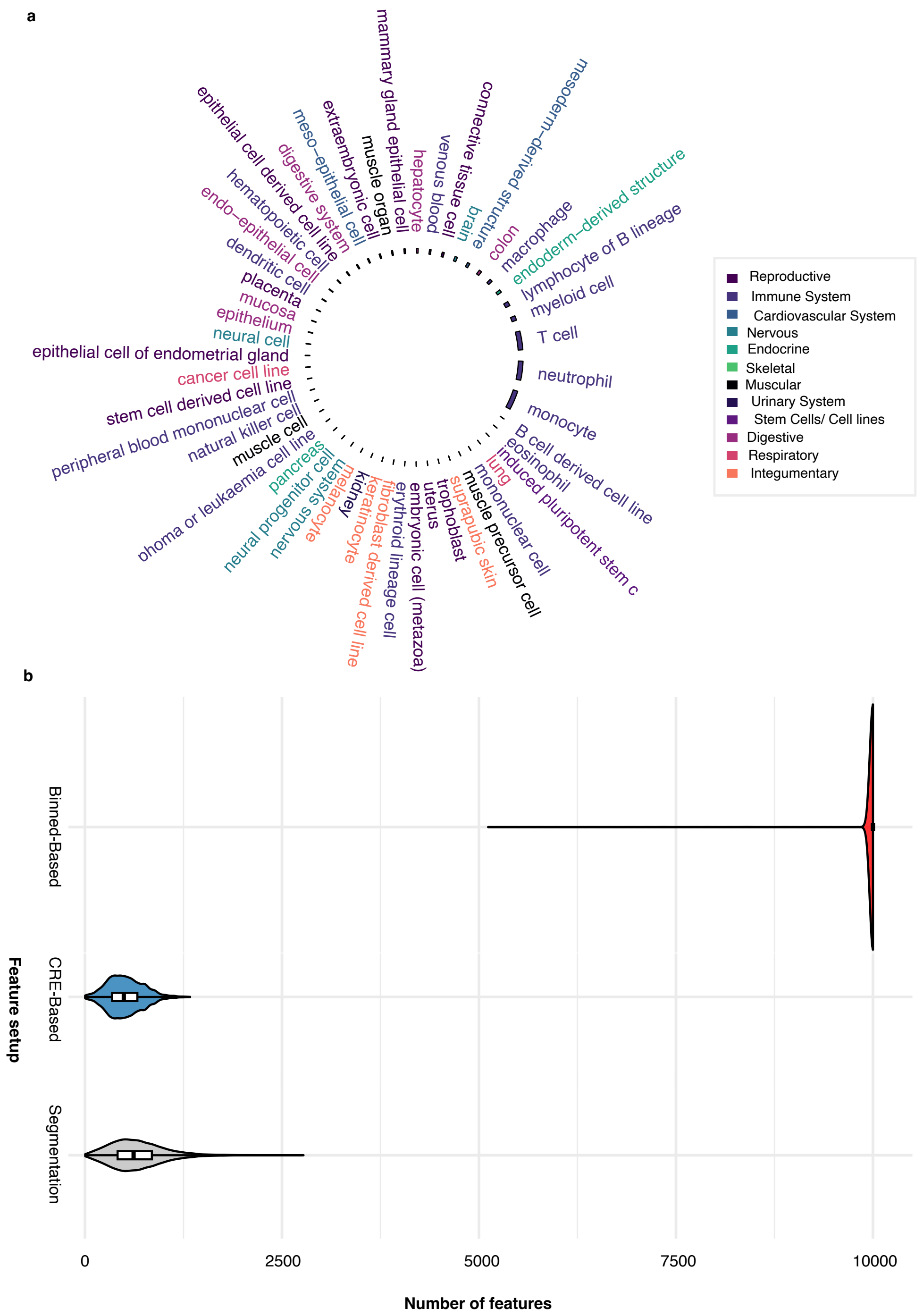


**Fig. 1**: **Number of samples per cell type and distribution of input features for different methods**. **(a)** Circos plot showing the number of samples of each cell type, colored by the higher-level lineage. The plot was generated with the circlize R package [1]. **(b)** Violin plots show distribution of input features for different feature setups.

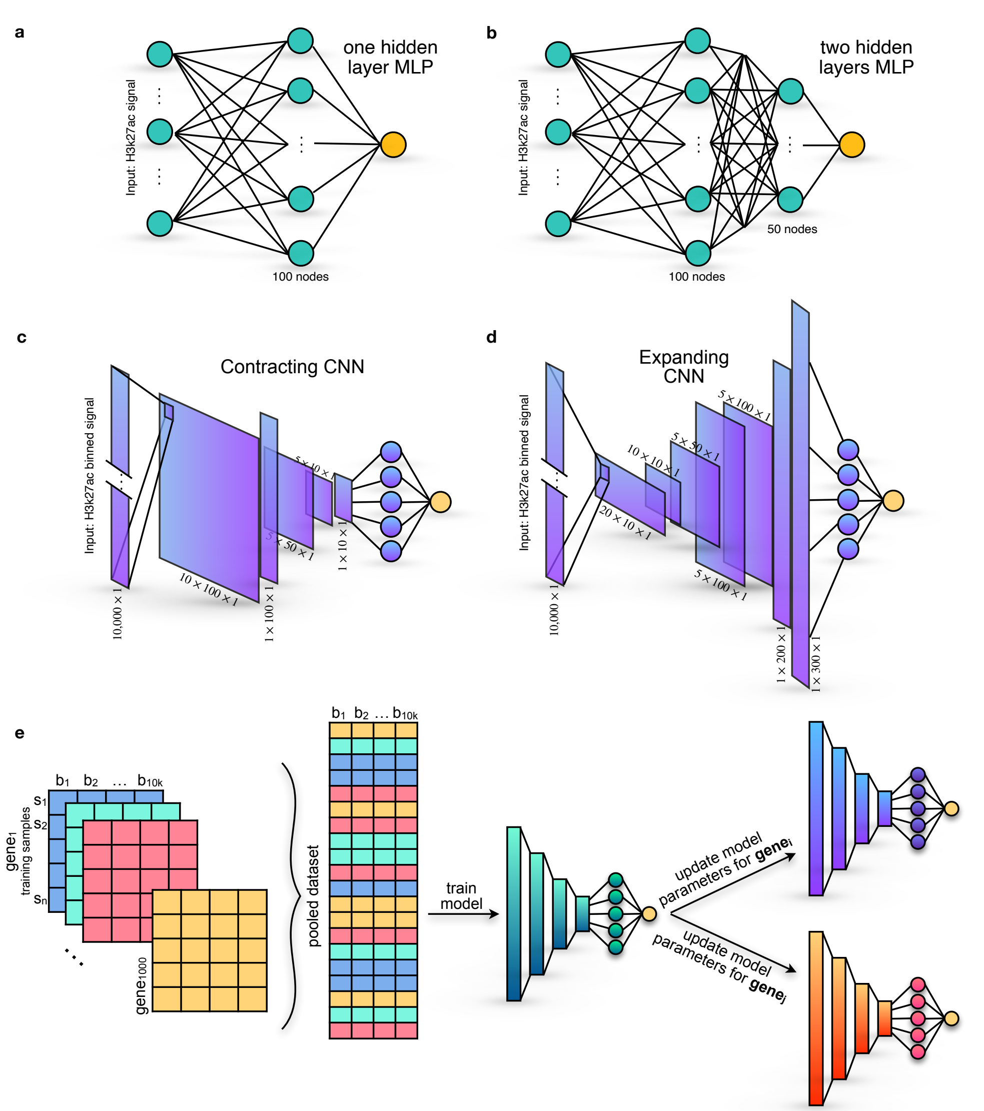


**Fig. 2**: **Architectures of the Multi-Layer Perceptron (MLP) and convolutional Neural Network (CNN) models and warm start strategy for CNN** **(a)** The MLP model with a single hidden layer consists of 100 nodes, fully connected to both the input layer and the output neuron. This model processes the H3K27ac signal as input and applies ReLU activation functions in the hidden layer, while the output layer utilizes a linear activation function for continuous predictions. The output of the model corresponds to the predicted RNA-seq expression count. **(b)** The MLP model with two hidden layers features 100 nodes in the first hidden layer and 50 nodes in the second hidden layer. Each layer is fully connected, and ReLU activation is applied to both hidden layers. The final output layer, similar to the single hidden layer model, employs a linear activation function. The output of the model also corresponds to the predicted RNA-seq expression count.**(c)** Architecture of the contracting convolutional neural network (CNN), where the kernel sizes progressively decrease from the input layer to the output layer. **(d)** Expanding CNN architecture, characterized by progressively larger kernel sizes from the input to the output layer. Both topologies include a fully connected dense layer with 5 hidden nodes linking the convolutional block to the output regression node which is designed to predict RNA expression counts. The ReLU activation function is applied to all layers except the first and last, where a sigmoidal activation function is used instead. Layer sizes are indicated in the figure. **(e)** A heuristic warm start approach was used for weight initialization of CNN models. Three datasets were created, each containing 1,000 randomly selected genes along with their corresponding biological samples. The contracting and expanding kernel CNNs were independently trained on these datasets, yielding optimized models for each subset. The learned parameters from these models were then used as initial weights to train CNN models for specific genes, improving convergence and stability. Genes located near chromosome borders, where a full 10,000-bin setup was unavailable, were trained using a cold start.

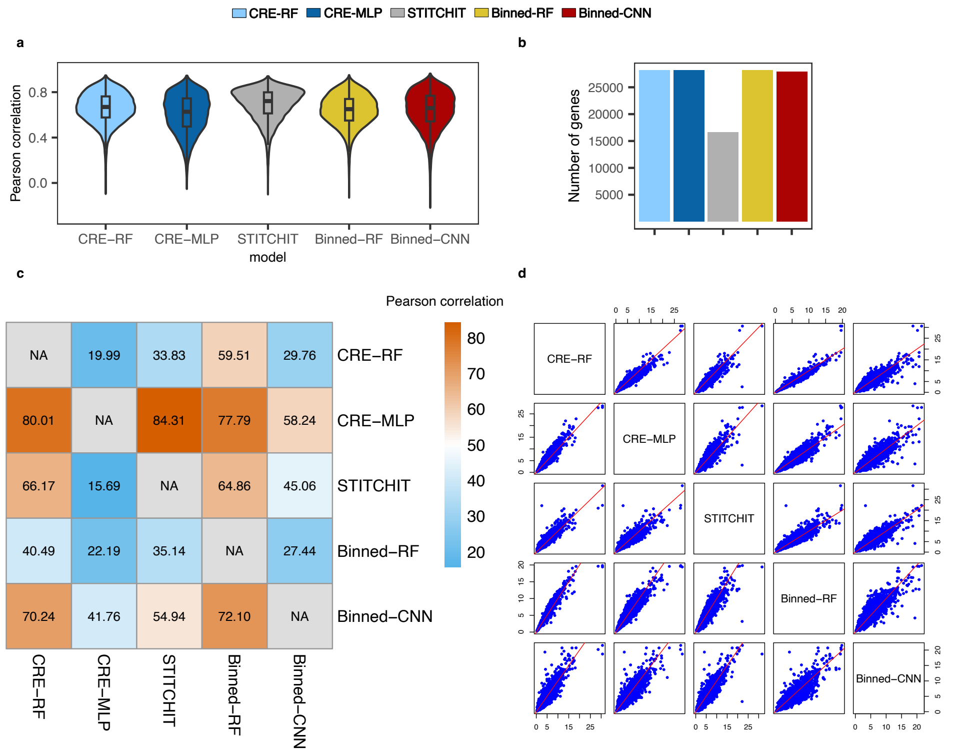


**Fig. 3**: **Performance assessment of the five machine learning approaches** **(a)** Violin plot of the Pearson correlation distributions per gene. Center lines of the included box plots are the median, boxlimits indicate the interquartile range (IQR), whiskers 1.5x IQR, outliers are removed. **(b)** Barplot of the number of the total learned models **(c)** Heatmap for the pairwise Pearson correlation comparison of the approaches, showing the percentage for which the method in the row outperforms the method in the column **(d)** MSE scatterplots for pairwise method comparison. Correlation and MSE are always estimated on the test set.

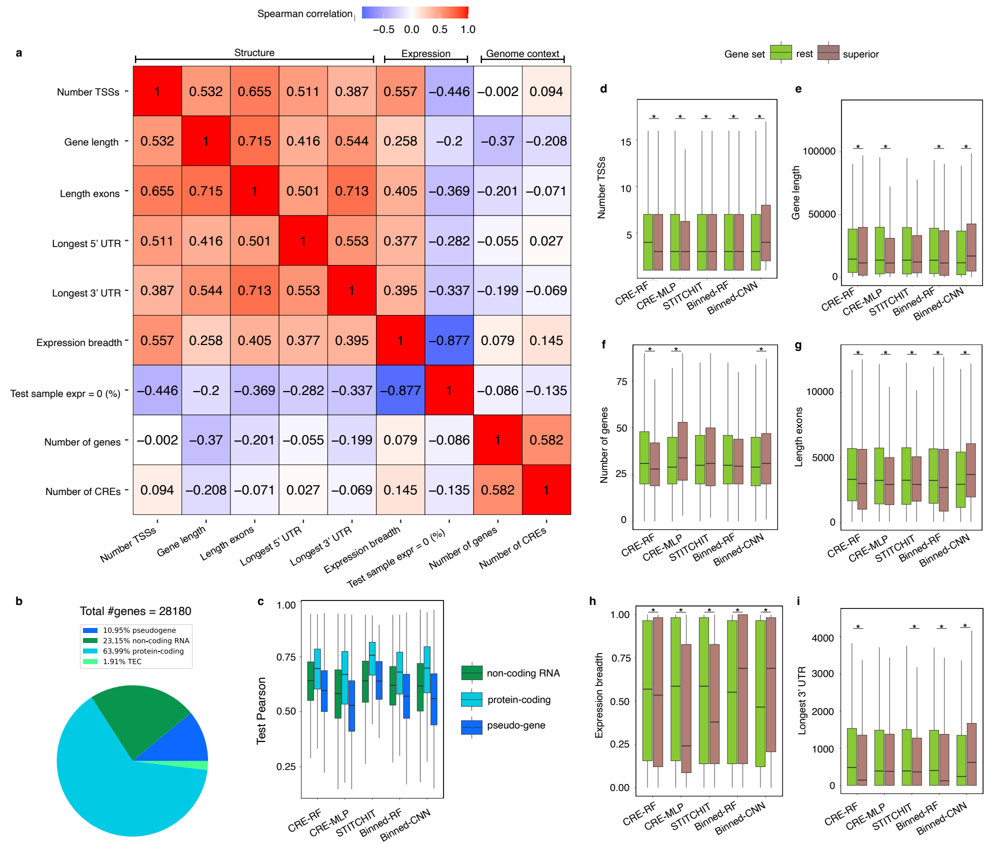


**Fig. 4**: **Investigation of gene characteristics that affect model performance.** **(a)** Spearman correlation heatmap for the gene descriptors. Descriptors that cluster together according to their correlation are grouped into three categories *structure*, *expression* and *genome context*. **(b)** Pie chart for the biotypes of the genes that have $\geq$ 2 expression variance and 90% non-zero values across samples in the RNA data. **(c)** Boxplot for the model performances (Pearson correlation) for each method on the subsets of genes that are non-coding, protein-coding and pseudogenes. **(d-i)** Boxplot for a gene descriptor showing the genes for which a method has the lowest MSE compared to the other methods (superior) and all the remaining genes (rest). The descriptors are **(d)** number of TSSs, **(e)** gene length, **(f)** number of other genes in the 1 MB gene window, **(g)** total exon length, **(h)** expression breadth, i.e. fraction of cell types/tissues (n = 58) in the entire dataset where the target gene is expressed (TPM $\geq$ 0.5), **(i)** length of the longest 3' UTR. For boxplots **(c-i)** the center line indicates the median, boxlimits correspond to IQR, whiskers to 1.5x IQR. An unpaired Mann-Whitney U Test was performed between the "superior" and "rest" gene sets for each gene descriptor and each method separately. A p-value $\leq$ 0.05 is indicated with an asterisk.Colours in **(b)** and **(c)** taken from the colorcet package based on Glasbey et al.. Pearson and MSE values were estimated on the test set.

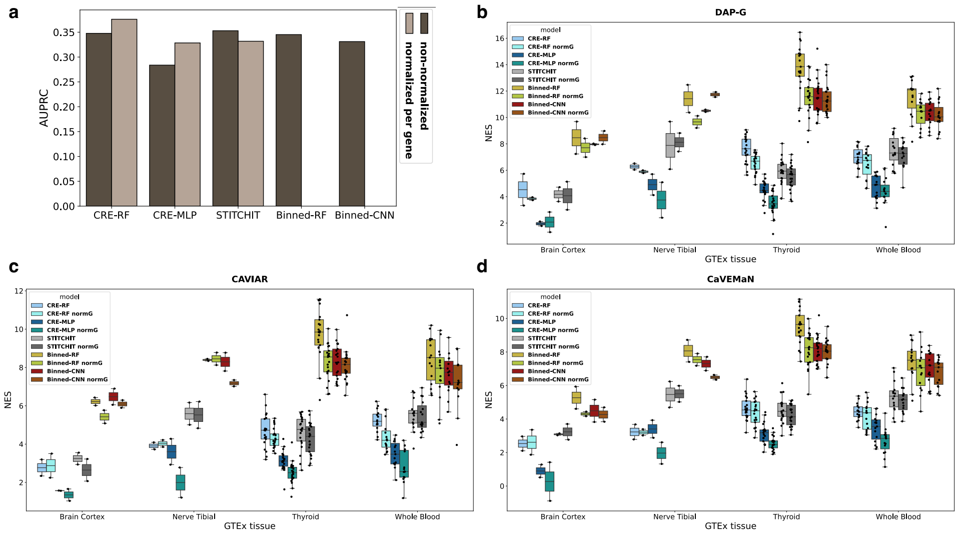


**Fig. 5**: **Model comparison using CRISPRi and eQTL data. (a)** Barplot with the area under the precision recall curve (AUPRC) of all models based on validated enhancer-gene interactions [3]. An *in silico* perturbation approach was implemented to assess the importance of an enhancer-gene interaction for a model. Two score calculations were tested: $ISP$ and $ISP_{normG}$. The AUPRC was calculated based on the set of interactions for which all models could produce a score for, namely 1,099 tested interactions out of which 198 were significant. For Binned-RF and Binned-CNN $ISP_{normG}$ was tested but is not shown, due to their feature space. Using the fused regions for normalization further reduced the set of interactions all model could produce a score for drastically, and normalizing across all bins was not computationally feasible. **(b-d)** Normalized enrichment score (NES) of the models for enhancer-gene interactions that are supported by eQTL-gene pairs from GTEx [4] (boxplot center line is the median, boxlimits show the inter-quartile range, whiskers up to 1.5x interquartile range). For all models both score calculations were tested. Here, $ISP_{normG}$ was possible for binned-RF and binned-CNN with the fused regions per gene, since the top 100,000 scored interactions were taken per model and not jointly. Shown are the eQTL-gene pairs fine-mapped with DAP-G [5] (b), CAVIAR [6] (c) and CaVEMaN [7] (d). The boxplots are formed by the NES of the EpiATLAS samples that were matched to the respective GTEx tissue.

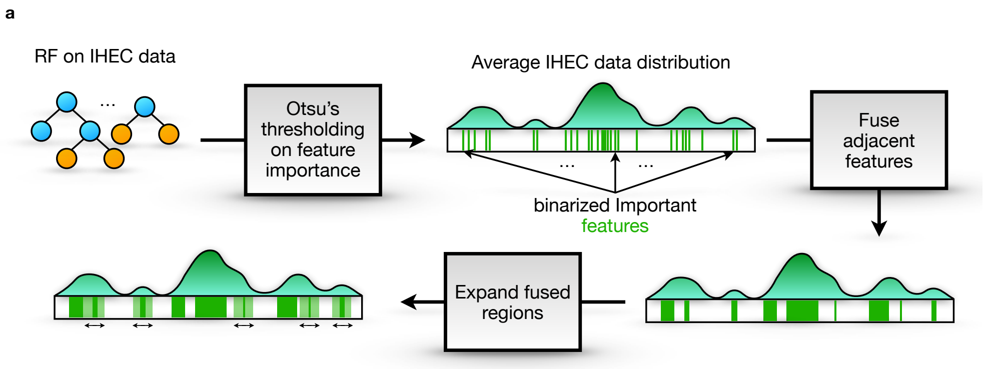


**Fig. 6**: **Schematic illustration of feature fusion in Binned-CNN models**. **(a)** Binned-RF are recruited to determine global feature importance for each gene. Then, Otsu's thresholding method is deployed to binarize feature importance values, classifying genomic bins as 'active' (1) or 'inactive' (0). To ensure meaningful feature selection, adjacent active bins are fused if they are separated by no more than five inactive bins, forming larger contiguous regions. Finally, these fused regions are further expanded to a minimum length of 1kb, through merging nearby (fused) regions, to form regions used for ISP analysis of the model.

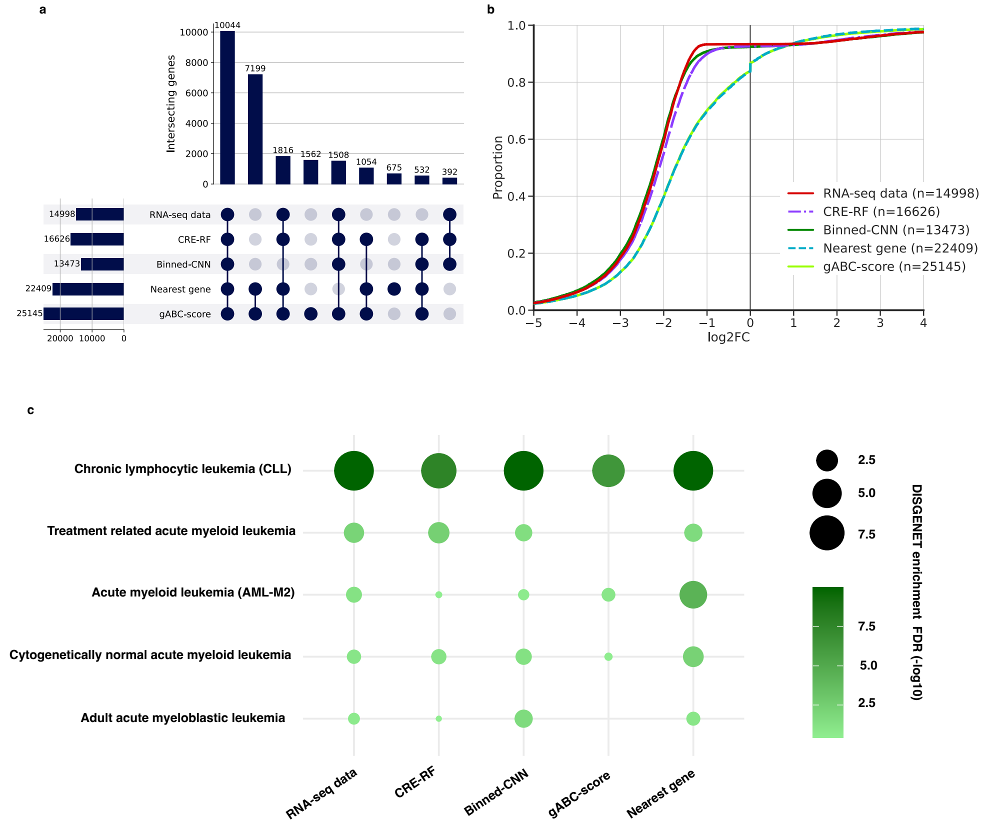


**Fig. 7**: **Comparison of genes found by the HAWAS-gene test and genes found by conventional approaches**. RNA-seq data: DEGs identified by DESeq2 [8] from the RNA-seq data. CRE-RF and Binned-CNN: Genes found with the HAWAS-gene test based on the CRE-RF and Binned-CNN models, respectively. Nearest gene: Genes that were closest to differentially acetylated regions identified with DiffBind [9], with a maximum distance of 100 kb. gABC-score: Genes that were associated to differentially acetylated regions according to the interactions predicted with the gABC-score [10]. All sets were limited to the 28,180 genes that were considered for model training. **(a)** UpSet plot showing the overlap of the gene sets, limited to the nine largest intersections [11]. **(b)** Cumulative distribution of the log2FC of the gene sets, as measured by the RNA-seq data. A positive log2FC indicates downregulation in CLL and vice versa. **(c)** Dot plot illustrating the enrichment of known leukemia genes (DisGeNET database [12]) among the gene sets (enrichment test -$\log_{10}$ FDR).

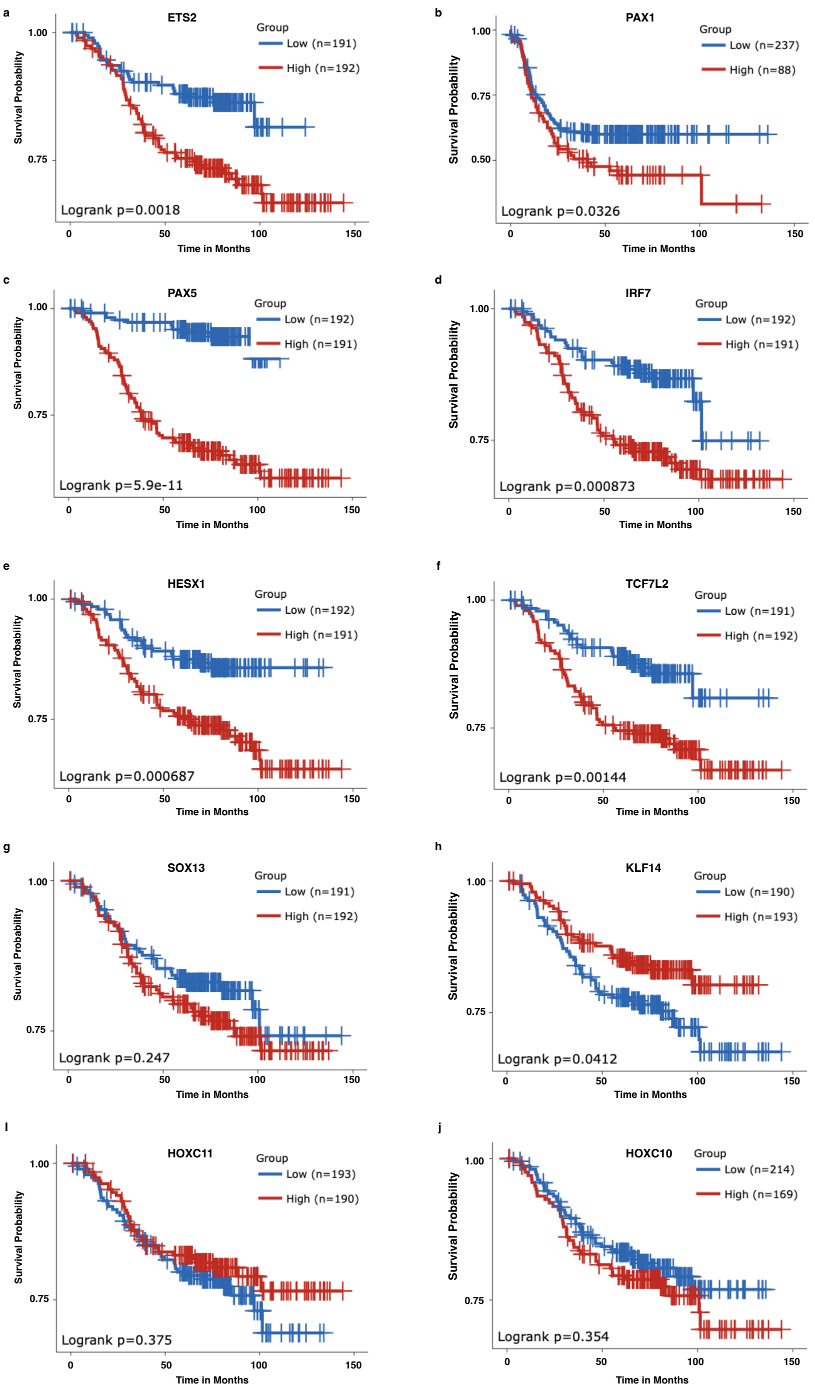


**Fig. 8**: **Kaplan-Meier survival analysis plots (a–j) for the top 10 TFs between control and CLL samples done with SurvivalGenie2.0 while using the TARGET-ALL-P2-Bone-Marrow dataset [13].** Each panel corresponds to a specific TF: **(a)** ETS2, **(b)** PAX1, **(c)** PAX5, **(d)** IRF7, **(e)** HESX1, **(f)** TCF7L2, **(g)** SOX13, **(h)** KLF14, **(i)** HOXC11, **(j)** HOXC10.

[4] Consortium, T.G., Aguet, F., Anand, S., Ardlie, K.G., Gabriel, S., Getz, G.A., Graubert, A., Hadley, K., Handsaker, R.E., Huang, K.H., Kashin, S., Li, X., MacArthur, D.G., Meier, S.R., Nedzel, J.L., Nguyen, D.T., Segrè, A.V., Todres, E., Balliu, B., Barbeira, A.N., Battle, A., Bonazzola, R., Brown, A., Brown, C.D., Castel, S.E., Conrad, D.F., Cotter, D.J., Cox, N., Das, S., Goede, O.M., Dermitzakis, E.T., Einson, J., Engelhardt, B.E., Eskin, E., Eulalio, T.Y., Ferraro, N.M., Flynn, E.D., Fresard, L., Gamazon, E.R., Garrido-Martín, D., Gay, N.R., Gloudemans, M.J., Guigó, R., Hame, A.R., He, Y., Hoffman, P.J., Hormozdiari, F., Hou, L., Im, H.K., Jo, B., Kasela, S., Kellis, M., Kim-Hellmuth, S., Kwong, A., Lappalainen, T., Li, X., Liang, Y., Mangul, S., Mohammadi, P., Montgomery, S.B., Muñoz-Aguirre, M., Nachun, D.C., Nobel, A.B., Oliva, M., Park, Y., Park, Y., Parsana, P., Rao, A.S., Reverter, F., Rouhana, J.M., Sabatti, C., Saha, A., Stephens, M., Stranger, B.E., Strober, B.J., Teran, N.A., Viñuela, A., Wang, G., Wen, X., Wright, F., Wucher, V., Zou, Y., Ferreira, P.G., Li, G., Melé, M., Yeger-Lotem, E., Barcus, M.E., Bradbury, D., Krubit, T., McLean, J.A., Qi, L., Robinson, K., Roche, N.V., Smith, A.M., Sobin, L., Tabor, D.E., Undale, A., Bridge, J., Brigham, L.E., Foster, B.A., Gillard, B.M., Hasz, R., Hunter, M., Johns, C., Johnson, M., Karasik, E., Kopen, G., Leinweber, W.F., McDonald, A., Moser, M.T., Myer, K., Ramsey, K.D., Roe, B., Shad, S., Thomas, J.A., Walters, G., Washington, M., Wheeler, J., Jewell, S.D., Rohrer, D.C., Valley, D.R., Davis, D.A., Mash, D.C., Branton, P.A., Barker, L.K., Gardiner, H.M., Mosavel, M., Siminoff, L.A., Flicek, P., Haeussler, M., Juettemann,

T., Kent, W.J., Lee, C.M., Powell, C.C., Rosenbloom, K.R., Ruffier, M., Sheppard, D., Taylor, K., Trevanion, S.J., Zerbino, D.R., Abell, N.S., Akey, J., Chen, L., Demanelis, K., Doherty, J.A., Feinberg, A.P., Hansen, K.D., Hickey, P.F., Jasmine, F., Jiang, L., Kaul, R., Kibriya, M.G., Li, J.B., Li, Q., Lin, S., Linder, S.E., Pierce, B.L., Rizzardi, L.F., Skol, A.D., Smith, K.S., Snyder, M., Stamatoyannopoulos, J., Tang, H., Wang, M., Carithers, L.J., Guan, P., Koester, S.E., Little, A.R., Moore, H.M., Nierras, C.R., Rao, A.K., Vaught, J.B., Volpi, S.: The GTEx consortium atlas of genetic regulatory effects across human tissues. Science **369**(6509), 1318–1330 (2020) https://doi.org/10.1126/science.aaz1776 https://www.science.org/doi/pdf/10.1126/science.aaz1776
